# Specialized Shh-sensing cell with unique cilia and basal body in the forebrain ventricular epithelium

**DOI:** 10.64898/2026.08.10.744053

**Authors:** Arantxa Cebrian-Silla, Fiona R Dale-Huang, Stephanie A Redmond, Carmen Elena Aragon Ortiz, Juan Morianos, Marcos Assis Nascimento, Zhenmeiyu Li, Cristina Guinto, Susana González- Granero, Ricardo Romero-Rodriguez, Cathryn R Cadwell, Vicente Herranz-Pérez, Jose Manuel Garcia-Verdugo, Arnold Kriegstein, Eric Huang, Arturo Alvarez-Buylla

## Abstract

Ependymal (E1) cells, with their tufts of ∼50 motile cilia, line the walls of the brain ventricles and help propel the cerebrospinal fluid (CSF). The CSF is rich in signaling molecules, but the cellular targets that detect these signals and their function remain unknown. Here, we describe a distinct population of ependymal cells (E2) in the forebrain of mice and humans, the majority having only 1 or 2 cilia. These cilia were motile, but unlike E1 cells’ cilia, their pattern of motility and high expression of Arl13b and Inpp5e suggest a sensory function. E2 cells were characterized by an enormous, donut-like basal body that contained an increased number and size of subdistal appendages. In mice, E2 cells were mostly born in the embryo, but completed their differentiation in juveniles and young adults; they were found at higher densities in regions of high CSF flow and neurogenesis. E2 cilia contained the G protein-coupled receptor Smoothened, which accumulated in their cilia upon exposure to Sonic Hedgehog (Shh). Together, these findings identify E2 cells as a novel CSF-sensing ependymal cell type and provide a cellular target for the CSF signaling.

## Introduction

The brain’s parenchyma is separated from the cerebrospinal fluid (CSF) in the ventricles by an epithelial lining of multiciliated ependymal (E1) cells^1,2^. These cells serve as a barrier and filter for CSF-derived factors, and, through the coordinated and constant beating of their cilia, they propel the CSF through the brain ventricles. The CSF was considered for many years as a simple brain mechanism for shock absorption and the removal of molecular waste^3^. However, more recent studies indicate that the CSF is a rich source of metabolites and signaling molecules, providing trophic and angiogenic factors, chemorepellents, small peptides, and their carrier proteins^4–7^. Therefore, the CSF represents a rich signaling milieu for a “private” circulatory system within the central nervous system. In this context, ependymal cells not only function as a selective filter, but contribute to homeostasis, the regulation of the neuroinflammatory response and aging^8–10^.

The forebrain ventricular epithelium also contains the thin apical processes of a population of astroglial (B1) cells that function as neural stem cells (NSCs) in the generation of neurons and oligodendrocytes in the adult brain^11–13^. Multiciliated ependymal cells (E1) contribute to the organization and regulation of this germinal activity in the ventricular-subventricular zone (V-SVZ)^14–16^. The small apical processes of B1 cells contain a single primary cilium and are surrounded by rosettes of E1 cells forming pinwheel structures. During the characterization of these pinwheels, a population of ependymal cells that apparently only had two cilia associated to two large donut-like basal bodies (E2 cells), was described^11^. E2-like cells have been also described in the third and fourth ventricles and in the central canal of the spinal cord, in rodents and humans^17–19^. However, the molecular and cellular features that define E2 cells as a distinct cell type, their functional significance, and their presence in the human lateral ventricles have yet to be determined.

Here, we combine molecular, ultrastructural, developmental, and functional analyses to define the identity and biological properties of mouse forebrain E2 cells. We found that E2 cells are not only biciliated, but are heterogenous, with most having one cilium. E2-like cells are also present in the walls of the adult human lateral ventricles. We identify the components responsible for the large donut-like basal body of E2 cells and show that their cilia exhibit a unique mode of linear motility distinct from that of E1 cells. The unique structure and motility of E2 cells cilia suggested that they may sample the CSF for molecular signals. Consistently, we show that E2 cells cilia respond to Sonic Hedgehog signaling. Together, these findings identify E2 cells as a novel sensory ependymal cell type and reveal a previously unrecognized sensory specialization within the ventricular epithelium.

## Results

### A novel population of forebrain ependymal cells with hybrid ciliary properties

Using confocal microscopy three-dimensional reconstructions, we determined that the basal bodies of E2 cells had a mean volume of 1.8 ± 0.5 um^3^ (n=19) about 45 times larger than those of neighboring E1 cells (0.045um^3^ ± 0.01, n=33 cells, Figure S1A). To characterize the molecular composition of E2 cell cilia, we stained wholemount preparations of the ventricular lateral wall with antibodies against ADP ribosylation factor like GTPase 13B (Arl13b), a protein essential for cilia formation and maintenance^20^. We combined this staining with γ-tubulin, and β-catenin to delineate the apical cell boundaries. Based on the presence of two large donut-like basal bodies and Arl13b staining we observed three subpopulations of E2 cells: 1) cells with no cilia, 2) cells with a single long cilium, and 3) cells with two long cilia (Figure 1A, B). In addition, we identified a few E2 cells with four large donut-like basal bodies and 0-4 cilia (Figure 1B). In young adults (P30), most E2 cells had either one cilium (55.1% ± 4.1) or two cilia (24.5% ± 4.5), while smaller proportions of E2 cells lacked cilia (10.0% ± 3.8) or had four basal bodies (10.4% ± 2.7) (n=6) (see below how these proportions change with age). Both uni- and biciliated E2 cells expressed markers of multiciliated ependymal cells (E1 cells) such as FoxJ1, S100b, Cd24, p73 and Vimentin, as well as Sox2 (Figure S1B). To study the morphology of E2 cells, we used P60 FoxJ1::CreERt2 in mice carrying a floxed allele Ai14 reporter mice. Ai14 expression was induced by the administration of low concentrations of tamoxifen to label isolated E cells. The cell body of E2 cells, for both uniciliated and biciliated cells, varied in morphology, from cuboidal to elongated and was smaller than that of E1 cells (Figure 1C and S1C-E). Some E2 cells’ somata were partially subependymal, with a smaller apical surface and part of the cytoplasm under neighboring E1 cells (Figure S1E, F). Interestingly, E2 cell morphology varied along the V-SVZ. Anteroventral E2 cells extended a long, thin basal process, whereas dorsal E2 cells frequently exhibited a shorter tangential process beneath neighboring E1 cells (Figure S1D-E). Thus, despite their morphological heterogeneity, E2 cells were consistently identified by their large, donut-like basal bodies.

**Figure 1:**
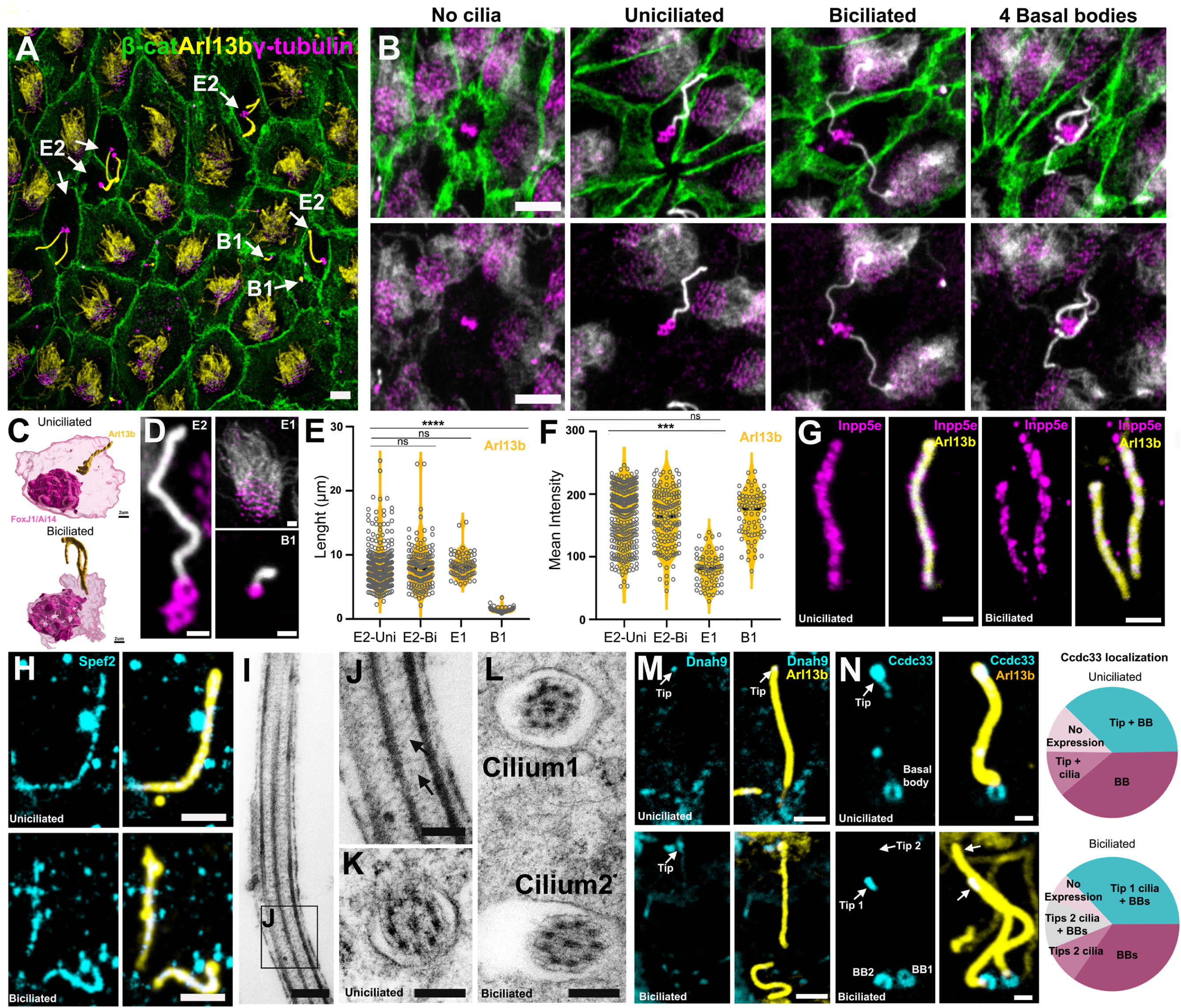
E2 cells cilia have properties of primary cilium and motile cilia (Related to Figure S1). **(A)** Confocal images of P30 whole mount of the lateral ventricular wall. E2 cells were identified by their large γ-tubulin donut-like basal bodies (magenta) and long Arl13b bright cilia (yellow). β-catenin (green) delineates cell borders. **(B)** High magnification images of E2 cells subtypes. **(C)** Three-dimensional reconstructions of uni- and bi-ciliated E2 cells based on *FoxJ1CreER:Ai14* and Arl13b expression. (**D**) Closeups of E2, B1, and E1 cell basal bodies (magenta) and cilia (white). (**E**) Quantifications of the length of E2, E1, and B1 cell cilia using Arl13b staining (uniciliated: 8.13 µm ± 0.48, 296 cells; biciliated: 8.04 µm ± 1.01, 72 cells, E1 cells 8.32 µm ± 1.21, 72 cells, B1 cells 1.40 µm ± 0.12, 72 cells n=4). (**F**) Quantifications of Arl13b mean intensity fluorescence in E2, E1, and B1 cell cilia (Uni E2: 160.19 ± 23.92, 296 cells; Bi E2: 159.65 ± 19.48, 72 cells vs E1: 80.78 ± 8.50, 72 cells, B1 cells: 166.91 ± 25.37, 72 cells n=4). **(G)** Inpp5e expression (magenta) in uni- and biciliated Arl13b E2 cells’ cilia. **(H)** Spef2 expression (Cyan) along Arl13b E2 cells cilia. **(I-L)** Electron micrographs from the cilia of uniciliated and bi-ciliated cells showing radial spokes (arrows) and a 9+2 central pair microtubule structure. **(M)** Confocal images of E2 cells’ cilia (Arl13b, yellow) showing Dnah9 expression (cyan) at the most distal tip. **(N)** Confocal images showing Ccdc33 localization (cyan) in E2 cell cilia and basal bodies, revealing heterogeneous distribution patterns. Pie charts summarize the proportions of Ccdc33 localization along the basal body-cilia axis. Uniciliated: 39% ±7.91 showed basal body-restricted labeling, 37% ± 8.25 showed Ccdc33 localization at both the basal body and distal ciliary tip, 11% ± 1.91 displayed signal along the ciliary shaft and tip, and 12% ±4.41 showed no detectable expression (100 cells, n = 3). Biciliated: 36% ± 4.65 both basal bodies with enrichment at the tip of only one cilium, 34% ± 10.61 basal body-restricted labeling, and 9% ±4.65 ciliary tips and basal bodies; the remaining cells exhibited either tip-restricted or undetectable expression (87 cells, n = 3). Test: one-way ANOVA. Statistical significance was defined as ∗∗*p* < 0.01, ∗∗∗*p* < 0.001, ∗∗∗∗*p* < 0.0001. Scale bars: 10µm (A),5µm (B, G, H), 2 µm (M), 1µm (D), 500nm (N), 200 nm (J-L), 100 nm (I).

To identify E2 cells cilia properties, we quantified the cilia length using Arl13b staining. We found that uni- and biciliated E2 cell cilia were similar in length (8.13 µm ± 0.48 and 8.04 µm ± 1.01, n = 4), comparable to those of E1 cells (8.32 µm ± 1.21), but much longer than the primary cilia of B1 cells (B1 cells 1.40 µm ± 0.12) (Figure 1D, E)^11^. Notably, E2 cells had higher Arl13b expression than E1 cells, at levels similar to those observed in B1 cells (Figure 1D, F). Arl13b interacts with Inpp5e, a ciliary phosphoinositide 5-phosphatase involved in ciliogenesis and ciliary function^21^. To assess Inpp5e expression in E2 cell cilia, we co-stained whole mounts for Inpp5e and Arl13b. Similar to Arl13b, Inpp5e was highly expressed in both uni- and biciliated E2 cell cilia (Figure 1G). Together, these results show that the majority of E2 cells have one or two long cilia with high expression of Arl13b and Inpp5e, proteins associated with primary cilium signaling.

Given that E2 cells share features with E1 cells, we next examined whether their cilia show properties of motile cilia. To distinguish between a primary (9+0) and motile (9+2) axonemal organization, we stained for sperm flagellar protein 2 (Spef2), a component associated with the central microtubule pair of motile cilia^22^. Co-staining with Arl13b showed Spef2 expression along the entire length of cilia in both uni- and biciliated E2 cells (Figure 1H), similar to E1 cilia and absent in the primary cilium of B1 cells. Transmission electron microscopy (TEM) reconstructions (n = 25 cells) confirmed that E2 cilia contain a central pair of microtubules (9+2 axoneme) and radial spokes (Figure 1I–L), hallmark features of motile cilia. We next examined two motile cilia-associated proteins: Dnah9, an axonemal dynein^23^, and Ccdc33, a regulator of ciliary organization and function in multiciliated cells^24^. In E2 cells, both Dnah9 and Ccdc33 localized to the distal ciliary tip, consistent with their localization in E1 cells (Figure 1M, N and S1G-I). Notably, Ccdc33 was also detected at basal bodies in both E1 and E2 cells, in contrast to other cell types^24^. Dnah9 localization in E2 cells was more restricted to the ciliary tip and exhibited lower fluorescence intensity than in E1 cells (Figure S1H). In contrast, Ccdc33 showed marked diversity in its subcellular localization across E2 cells. Labeling was detected at basal bodies, ciliary tips, and occasionally along the ciliary shaft, with distinct combinations observed in both uni- and biciliated cells, suggesting dynamic regulation of Ccdc33 within E2 cell cilia (Figure 1N). These results reveal motile cilia features in E2 cells, with substantial heterogeneity in Ccdc33 localization, potentially reflecting differences in ciliary maturation or functional state.

Together, these results identify E2 cells as a distinct ependymal cell type defined by large, donut-like basal bodies and morphological heterogeneity, with most cells bearing one or two cilia that combine features of primary and motile cilia.

### E2 cells are present in walls of the lateral ventricle in the human brain

We next asked whether E2 cells are present in the human lateral ventricles. We stained whole-mount preparations of the anterior lateral ventricular wall from human postmortem samples with antibodies against Arl13b, γ-tubulin and β-catenin (Table1). Arl13b staining revealed ependymal cells with one or two long cilia (Figure S1M). These cilia were similar in length to those of neighboring multiciliated E1 cells and to mouse E2 cell cilia (Uni: 8.15 ± 0.54 µm; Bi: 8.84 ± 0.53 µm; E1: 9.82 ± 0.52 µm; n = 3) (Figure S1L). Consistent with our quantifications in mice, the cilia of human uni- and biciliated cells showed higher Arl13b fluorescence intensity than neighboring E1 cilia (Uni: 152.590 ± 30.39; Bi: 140.218 ± 48.60; E1: 79.624 ± 24.47; n = 3) (Figure S1L, M). As previously reported γ-tubulin did not reliably detect basal bodies in postmortem samples^17^. We therefore used pericentrin as an alternative basal body marker and found ependymal cells with small apical surfaces with large donut-like basal bodies (Figure S1N). These results show that the human lateral ventricles contain E2 cells similar to those in mice, suggesting a conserved function in mammals’ lateral ventricles including humans.

### Large basal bodies in E2 cells correspond to greatly expanded subdistal appendages

A defining property of E2 cells is their large donut-like γ-tubulin^+^ basal body. Although these basal bodies are greatly expanded compared to previously described basal bodies^25–27^, the specific components underlying this unique enlarged architecture remain unclear. To understand E2 cell basal body organization, we performed (en face and transverse) TEM reconstructions of 70 nm serial sections from the walls of the mice lateral ventricles. The basal bodies of 7 uni- and 4 biciliated E2 cells were serially reconstructed. E2 cell centrioles were similar in length to those of E1 cells (E2: ∼350 nm; E1: ∼325 nm) and were anchored to the apical plasma membrane via distal appendages (transition fibers), which were structurally comparable to those observed in E1 cells, B1 NSCs, and other cell types (Figure S2A-C)^28,29^. However, directly beneath these distal appendages, large electron-dense substructures extended 250 to 650 nm from the centriolar wall (Figure 2A-E). In the 7 uniciliated cells, and in one of the two cilia of the 4 biciliated cells, the basal body displayed a radial pattern of nine spokes extending from the centriole (Figures 2A, D, S2D, E). Three-dimensional reconstruction revealed a progressive rotational shift of the radial spokes from the apical to the basal region of the basal body, resulting in a partial helical arrangement (Figures 2B). Anchored to the ends of each of these structures, we found microtubules extending radially along the apical domain of the cell (Figure 2F). This organization was further confirmed by immunofluorescence staining for α-tubulin in combination with γ-tubulin and β-catenin (Figure 2G, H). Their donut-like radial arrangement, position along the centriole, and connection to the microtubule network indicate that the distinctive basal bodies in E2 cells correspond to greatly enlarged subdistal appendages (sDAPs)^30,31^. The number and distribution of these sDAPs also allowed us to distinguish the mother and daughter centrioles within each basal body pair.

**Figure 2.**
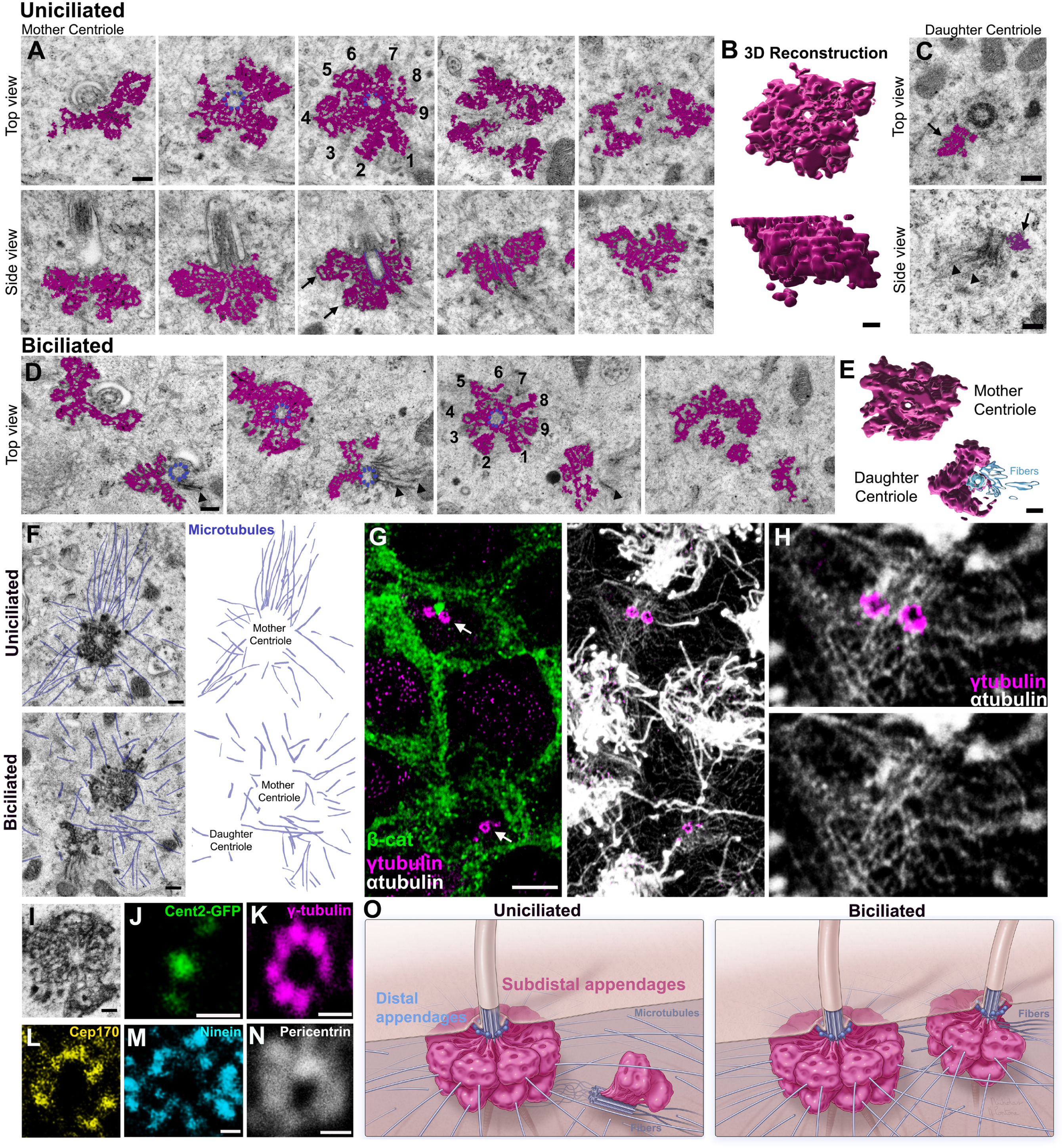
E2 cells’ basal bodies show unique subdistal appendages (Related Figure S2). (**A**) TEM ultramicrographs of serial sections of uniciliated E2 cells basal bodies (Sections spaced 140nm, see full reconstruction in Fig. S2). Top and side view of uniciliated E2 cells mother centriole showing the centriolar wall (colored in blue), associated with nine large, electron-dense subdistal appendages (sDAPs) colored in magenta. (**B**) Three-dimensional reconstruction of a uniciliated E2 cell mother centriole. (**C**) TEM micrographs of uniciliated E2 cells daughter centrioles in A, B, respectively, extending a single sDAP (magenta, arrows) and fibers (arrow heads). (**D**) TEM serial micrographs of biciliated E2 cell basal bodies. Note that the mother centriole shows sDAPs with 9 radial spokes, while the daughter centriole shows fewer sDAPs extending from three triplets of microtubules. Similar to uniciliated cells, the daughter centriole also shows fibers (arrowheads). (**E**) Three-dimensional reconstruction of a biciliated E2 cell basal bodies. (**F**) Superimposed drawings of microtubules (blue) from TEM serial reconstructions (910μm analyzed) showing that microtubules arrange around E2 cells’ basal bodies in a radial distribution. (**G, H**) Confocal images of E2 cells reveal a microtubule (α-tubulin) network extending radially from the basal bodies (γ- tubulin) to the cell borders (β-catenin). (**I**) High magnification TEM micrograph of a mother centriole extending sDAPs. (**J-N**) Confocal images of E2 cells basal bodies showing the protein distribution of subdistal appendages from the centriolar wall (Cent2:GFP) to the periphery (Pericentrin). Note that γ-tubulin, Cep170 and Ninein form concentric rings around the centriole. (**O**) Model of uni-ciliated and biciliated E2 cell basal bodies organization, illustrating distal and sDAPs. Scale bars: 500nm (J, K, L, M, N), 200nm (A, B, C and D), 250 (F, G, H, I).

Accordingly, the centriole bearing a complete set of nine sDAP spokes was identified as the mother centriole in both uni- and bi-ciliated E2 cells. In uniciliated cells, the mother centriole was linked to the daughter centriole via inter-centriolar fibers. In six of the seven reconstructed uniciliated cells, the daughter centriole displayed a single spoke within the sDAP region, and its proximal end extended bundles of fibers and ciliary rootlets (Figure 2C). In addition, all reconstructed uniciliated cells exhibited an electron-dense appendage close to the mother centriole but not connected to the centriolar wall (Figure S2D).

In biciliated cells, reconstructions revealed that the daughter centriole was the basal body of the second cilium. Similar to uniciliated cells, the daughter centriole of biciliated cells had sDAPs. However, these structures were more elaborate than those in uniciliated cells, showing multiple spokes that partially wrapped around the centriolar wall in a horseshoe-like configuration (Figure 2D, E). In addition to sDAPs, both uni- and biciliated cells showed pericentriolar satellites and multivesicular bodies, both closely associated with the microtubules extending from the sDAPs (Figure S2D, E), suggesting active vesicular trafficking.

To identify the ultrastructural correlate of the organization of E2 cells’ basal bodies with the previously described donut-like γ-tubulin^+^ structure^11^, we performed γ-tubulin immunogold staining. Gold particles localized to the electron-dense sDAPs, whereas no labeling was detected along the centriole (Figure S2F). To confirm this finding, we performed immunofluorescence for γ-tubulin and Arl13b in Centrin2: EGFP reporter mice, which label centrioles. Consistently, γ-tubulin formed a donut-like or horseshoe-like shape around Centrin2 (Figure 2J, K). Notably, in about 80% of the biciliated cells (65 cells, n=3), one cilium was associated with a γ-tubulin^+^ donut-like structure, while the second cilium corresponded to a horseshoe-like γ-tubulin structure (Figure S2G). In biciliated cells, the number of sDAPs allowed to distinguish the mother and daughter centrioles, consistent with the TEM reconstructions above. Our TEM analysis revealed that enlarged sDAPs in E2 cells serve as microtubule anchoring sites. Cep170 and Ninein are linked to γ-tubulin in the periphery of sDAP and contribute to microtubule nucleation and anchorage^32^. In E2 cells we found that Cep170 and Ninein were localized at the periphery of γ-tubulin (Figure 2K-M). Consistent with their role in microtubule organization, Pericentrin, a key scaffolding protein that anchors centrosomal components^33^, was also detected surrounding the basal bodies (Figure 2N). Altogether, these results show that E2 cells’ basal bodies are greatly enlarged sDAPs that anchor microtubules associated with multivesicular bodies and satellites (Figure 2O), suggesting enhanced microtubule anchorage and vesicular trafficking.

### E2 Cells have motile cilia with unique stroke patterns

The above results show that E2 cells cilia show features of motile cilia. However, whether these cilia actively move, the type of motility they display, and whether their movement is directionally polarized remains unknown. To address these questions, we performed live imaging recording of whole mounts from Arl13b-mCherry; Centrin-GFP mice to visualize E2 cells’ cilia and basal bodies, respectively^34^. Whole mounts of P30 mice were maintained in artificial CSF at room temperature under a confocal microscope. E2 cells were identified by their brightly labeled mCherry cilia and GFP^+^ centrioles (Figure 3A, B). We analyzed ciliary movement in 9 uniciliated and 7 biciliated E2 cells. The overall direction of CSF flow was determined by the polarized beating of neighboring E1 cilia^15,35^. We found that E2 cilia were motile with the proximal segment of the ciliary shaft (2-3 µm from the basal body) exhibiting a linear back and forth movement with a range of motion of 70.4° (±19.6°) (Figure 3C-J, Figure S3A, B and Movies S1-2). The ciliary tip generally tracked the angular movement of the base but moved more loosely (Figure 3C-J). This type of motility differs from the stereotypical movement of E1 cilia^36^. E2 cilia did not appear to have distinct effective and recovery strokes, with the tip of their cilia remaining extended throughout the entire cycle. We also observed that E2 cilia did not always complete full strokes; sometimes they partially completed the displacement in one direction before changing direction (Movie S2, Figure 3K). The whiplash-like movement of the distal portion of E2 cilia resulted in deviation from the plane swept by the proximal portion, resulting in the tips covering a larger oval-shaped area (Figure 3E, I).

**Figure 3:**
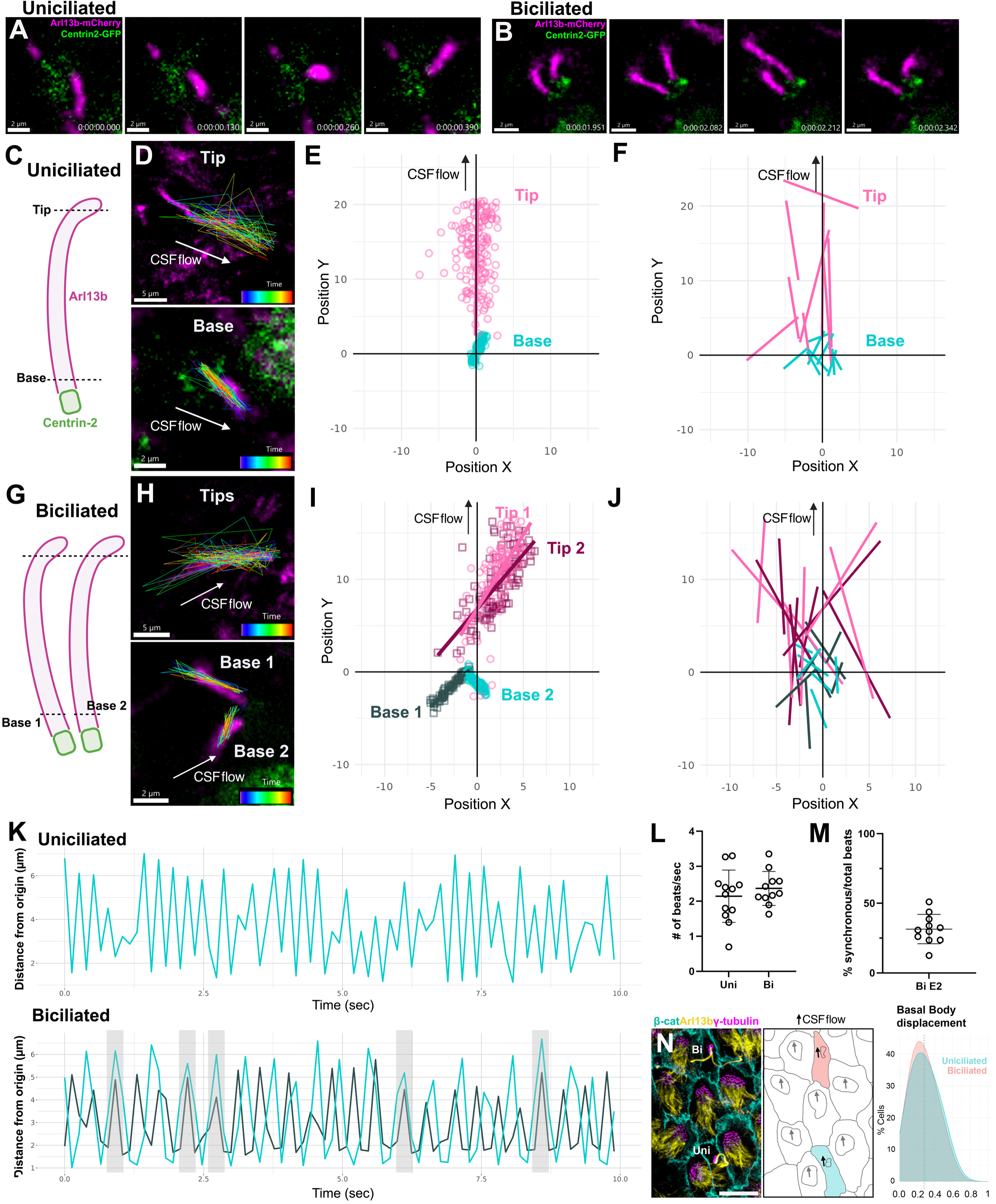
**E2 cell cilia are motile and have unique linear stroke patterns (Related to Figure S3 and movies, M1 and M2)**. **(A, B)** Frame-by-frame still images of a stroke in a uni-ciliated (A) and a bi-ciliated (B) E2 cells. Timestamps show frames each 13 second recording. **(C, D)** Frame-by-frame tracking of the position of the ciliary tip (top panel) and base segment (bottom panel) of a uniciliated E2 cell during the recording. CSF flow (inferred from neighboring E1 cells’s beating direction) marked with a white arrow. Colored bar depicting the passage of time in the tracking lines. **(E)** Scattered plot of the position (frame-by-frame) of the tip and the base segments of the cilium depicted in **D**. **(F)** Plot showing the regression line of frame-by-frame positions of the tips and of the base segments of 9 uniciliated E2 cells. **(G, H)** Frame-by-frame tracking of the position of the tip (top panel) and proximal segment (bottom panel) of a biciliated E2 cell. **(I)** Scattered plot of the position (frame-by-frame) of the tip and the base segments of both cilia depicted in **H**. **(J)** Plot showing the regression lines of the tips and base segments of 7 biciliated E2 cells. **(K)** Frequency diagram of the position of the base segment of the cilia in a uniciliated cell (top panel) and bi-ciliated cells (bottom panel). For the bi-ciliated cell, the shaded gray rectangles highlight synchronous strokes. **(L)** Quantification of the number of beats/secs observed at the proximal segment of uniciliated and biciliated E2 cells (n=12 and 11 respectively). **(M)** Quantification of the percentage of synchronous beats out of the total number of beats performed by biciliated E2 cell cilia. (**C**) Confocal images from the AV region of the lateral wall wholemounts stained for Arl13b (yellow), β-catenin (cyan) and γ-tubulin (magenta). (**D**) Traces of the apical surface and basal bodies of E1 and E2 cells in (A) showing basal bodies displacement. (**E**) Histogram of the basal bodies position of E2 cells relative to the apical surface. 0 represents the center mass and 100 the cell border. Average distance for uni ciliated (0.286 ± 0.01, 154 cells, n=3) cells and biciliated (0.272 ± 0.05, 83 cells, n=3) cells. Scale bars, 2µm (A, B, D, H), 10µm (N).

The overall direction of ciliary beating in uni- and biciliated E2 cells was similar, but not identical to that of neighboring cilia of E1 cells. In some cells, the beating direction of E2 cells was almost perpendicular to that of neighboring E1 cells (Figure 3A, B). The cilia of uniciliated E2 cells beat with an average frequency of 2.1 Hz (±0.7 Hz) at room temperature, and biciliated E2 cells at an average of 2.4 Hz (±0.5 Hz) (Figure 3K, L). This is much slower than the beating frequency observed in E1 cells at room temperature (30-50 Hz)^37,38^. Interestingly, in contrast to the synchronous beating observed in E1 cilia, the two cilia of biciliated E2 cells beat independently at different frequencies. The slight differences in beating frequency resulted in synchronous strokes in about one in three cycles (31.5% (±10.5)) for the duration of the recordings (Figure 3M). We also recorded an E2 cell with 4 basal bodies and 4 cilia, where each cilium beat independently.

The above results indicate that E2 cilia are motile. Hinged at the basal body, the cilia in E2 cells move back and forth in a linear fashion. The direction of movement varied but was largely similar to the movement of cilia in neighboring E1 cells. Furthermore, the angles of beating and frequency of individual cilia in E2 cells with 2 or more cilia were not correlated.

### E2 cells’ cilia display translational polarity

The above findings show that E2 cells’ cilia are motile and their motility is polarized similarly to that of E1 cells. In E1 cells, the orientation of the sDAP (basal foot) determines the rotational polarity of each cilium and directs the CSF flow^35,39,40^. However, in E2 cells, the mother centriole, despite its greatly expanded radially organized sDAPs, did not show evident rotational polarity (Figure 2O). In addition to rotational polarity, E1 cells display translational polarity through the asymmetric positioning of their ciliary tufts on the apical surface^35,41,42^. We investigated whether E2 cell cilia emerged from the center of the apical surface, and if displaced, in what direction compared to the polarization of neighboring E1 cells. We analyzed 154 uniciliated and 83 biciliated E2 cells in whole-mount preparations from P60 mice. In both uni- and biciliated E2 cells, the basal bodies were asymmetrically positioned, with a normalized displacement of 0.286 ± 0.01 and 0.272 ± 0.05, respectively (n = 3), from the center of the apical surface (Figure 3N). However, this displacement was smaller than that reported for E1 cells (0.4)^11^. Interestingly, we found that in more than half of E2 cells (Uni: 67.09±11.3 and Bi: 62.27±14, n=3) the mother centriole was positioned at the front of the pair in the direction of the basal body displacement (the direction of translational polarity) (Figure S3D). These findings indicate that, although less pronounced than in E1 cells, E2 cells exhibit translational polarity oriented in the direction of CSF flow.

### E2 cells are enriched in distinct domains and show regional specification

As shown above, γ-tubulin, Arl13b, and β-catenin stainings revealed four distinct subtypes of E2 cells: 1) uniciliated cells, 2) biciliated cells, 3) cells with no cilia, and 4) cells with four donut basal bodies and a variable number of cilia (1 to 4) (Figure 4A). We mapped the locations of these 4 subtypes in a whole mount at P60 (2,631 E2 cells counted). E2 cells were observed throughout most of the lateral wall, with the highest concentrations in the anterior-ventral (AV) and the posterior-dorsal (PD) regions, the dorsal corridor just ventral to the corpus callosum (Figures 4A and S4A). Across the whole mount, 51.5% of E2 cells were uniciliated, 23.4% were biciliated, 13.0% had no cilia, and 12.1% had 4 basal bodies and a variable number of cilia.

**Figure 4.**
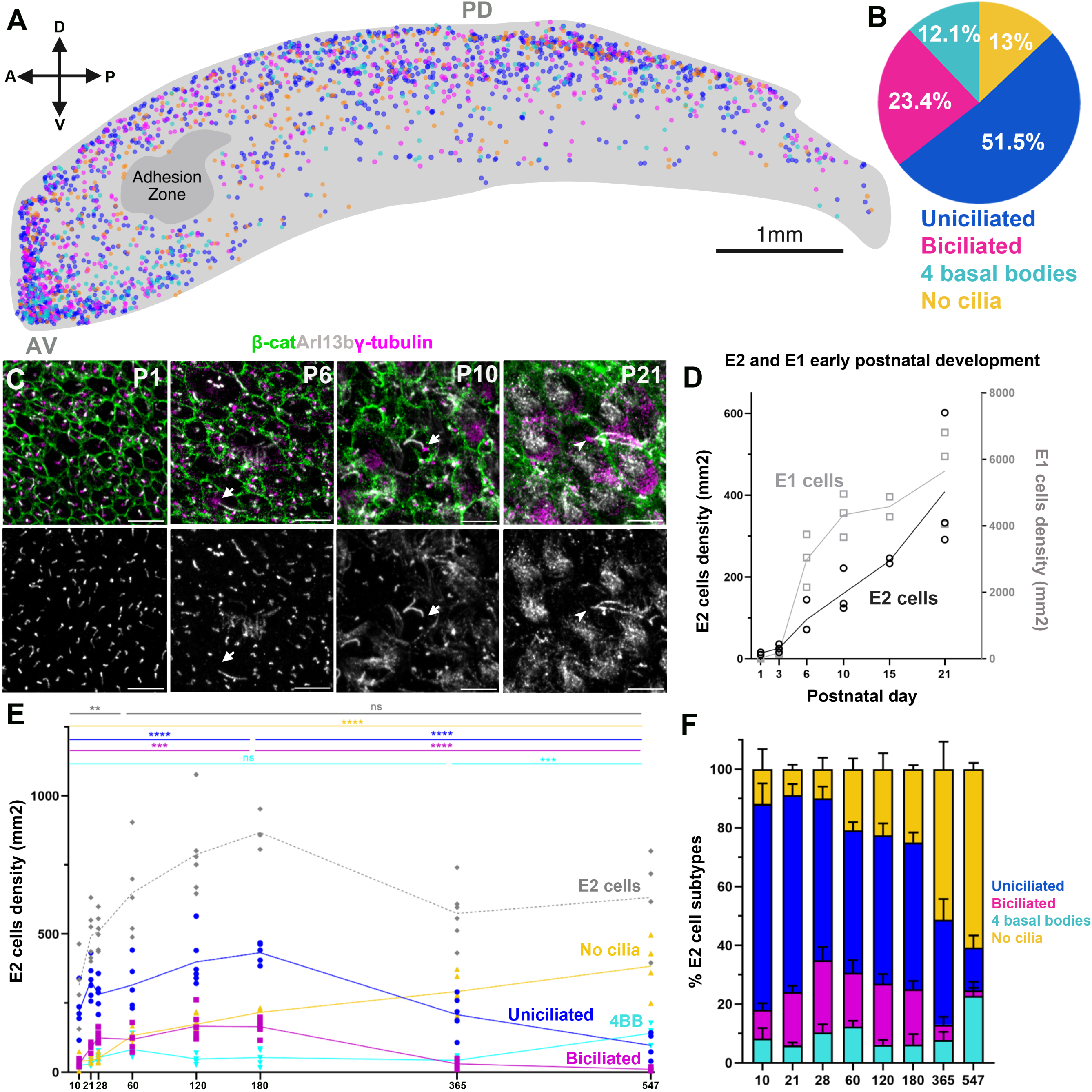
E2 cells spatial and temporal distribution (Related to Figure S4). **A)** Wholemount map of the distribution of E2 cell subtypes at P60. The compass indicates anterior (A), posterior (P), ventral (V), and dorsal (D) directions with the anterior-ventral (AV) and posterior-dorsal (PD) regions labeled. **(B)** Pie chart showing the proportions of E2 cell subtypes in the whole mount in A. **C)** Confocal images of the anteroventral most corner of the lateral wall at P1, P6, P10 and P21 stained with γ-tubulin (magenta), b-catenin (green), and Arl13b (white). Arrows indicate uniciliated E2 cells, arrowhead indicates a biciliated E2 cell with two long Arl13b bright cilia. (**D)** Density graph of E2 and E1 cells appearance after birth. The number of E2 cells duplicated by P3 (25.6 cells/mm^2^ ± s.d. 10.1) and continued to increase at P6 (95.9 ±42.1 cells/mm^2^), P10, (160.1 cells/mm^2^ (± s.d. 42.1), P15 (271.6 cells/mm^2^ (± s.d. 54.3), and P21 (408.6 cells/mm^2^ (± s.d. 168.6). **(E)** Quantifications of E2 cells subtypes over time in whole mounts (*n* = 6, 3 females and 3 males/time point, except P545 3 females and 1 male). E2 cells significantly increased from P10 and P60 (314.7 to 649.1 E2 cells/mm^2^). Both uni- and biciliated cells significantly increased from P10 to P180 (Uni: from 219.8 to 432.8 cells/mm^2^; Bi: from 29.7 to 164.5 cells/mm^2^), but then their numbers decreased between P180 to P547 (Uni: from 432.8 to 97.0 cells/mm^2^; Bi: from 164.5 to 11.38 cells/mm^2^). In contrast, E2 cells with no cilia increased with age (P10 to P547 from 40.10 to 383.9 cells/mm^2^). **(F)** Barplot showing the changes in the proportion of E2 cells subtypes over time. Test: one-way ANOVA with Tukey’s multiple comparisons. Statistical significance was defined as _∗_*p* ≤ 0.05, _∗∗_*p* < 0.01, _∗∗∗_*p* < 0.001, _∗∗∗∗_*p* < 0.0001. C: Scale bar, 5mm ( C).

NSCs in the walls of the lateral ventricle are regionally specified^43–46^. Differences in gene expression have consistently been noted between the dorsal and ventral V-SVZ, with the ventral domain characterized by the expression of Crym^43,47^. To determine whether E2 cells also show regional specification, we stained whole mounts of the lateral wall for Crym, in combination with β-catenin and Arl13b to identify E2 cells. We found that E2 cells in the PD region were Crym^-^, while 72.3% (± 3.8) of uniciliated and 37.2% (± 2.1) of biciliated cells in the AV were Crym^+^ (Figure S4C, n=3, cells=394). A subset of uniciliated E2 cells in the AV region (14.0% ± 2.1) showed particularly high levels of Crym expression (Figure S4C). The presence of Crym^+^ and negative E2 cells in the ventral V-SVZ and the protein level were confirmed using RNAscope *in situ* hybridization. Only a subpopulation of E2 cells in the ventral region expressed *Crym* (Figure S4D). These results suggest that E2 cells are regionally specified, but heterogeneous. In the dorsal regions, all E2 cells were Crym^-^, whereas a subpopulation in the ventral E2 cells expressed these markers associated with the regional specification of this epithelium.

### Developmental appearance of E2 cells and changes in their numbers with age

In mice, the ventricular surface of the lateral ventricles undergoes a dramatic remodeling during the first postnatal week, with most of the E1 cells maturing along a caudo-rostral and ventro-dorsal gradient between P0 and P10^48,49^. To determine when E2 cells first emerge, we analyzed whole-mount preparations of the lateral ventricles from P1, P3, P6, P10, P15, and P21 mice (n=3/age). E2 cells were identified by the presence of donut-like γ-tubulin_⁺_ basal bodies and Arl13b_⁺_ cilia. A few E2 cells (13.8 ± 3.3 cells/mm²) were observed at P1; at this age most cells lining the ventricular surface are radial glia characterized by small apical surfaces and short primary cilia (Figure 4C)^50^. By P3, the number of E2 cells duplicated (25.6 cells/mm^2^ ± s.d. 10.1) and continued to increase up to P21 (408.6 cells/mm^2^ (± s.d. 168.6). These findings indicate that, similar to E1 cells, E2 cells undergo postnatal differentiation. However, in contrast to E1 cells, which exhibit a sharp increase in number between P3 and P10, the increase in E2 cell number was more gradual (Figure 4D).

To further characterize age-related changes in the E2 cell population, we analyzed the AV and PD regions of whole-mount preparations at P10, P21, P28, P60, P120, P180, P365, and P547 (n=3/age/sex/region) (Figure 4E). In contrast to E1 cells, the overall density of E2 cells continued to increase between P10 and P60 from 314.7 to 649.1 E2 cells/mm^2^ (p-value=0.002). Consistent with the E2 cell map described above at P60, uniciliated cells were the most abundant E2 cell subtype in young mice, followed by biciliated cells, cells lacking cilia, and E2 cells with four basal bodies (Figure 4E, F).

However, the relative abundance of these E2 cell subtypes shifted markedly with age. The densities of both uni- and biciliated cells significantly increased from P10 to P180 (Uni: from 219.8 to 432.8 cells/mm^2^, p-value: <0.0001; Bi: from 29.7 to 164.5 cells/mm^2^, p-value: 0.0001) but subsequently declined between P180 and P547 (Uni: from 432.8 to 97.0 cells/mm^2^, p-value: <0.0001; Bi: from 164.5 to 11.38 cells/mm^2^, p-value: <0.0001). By contrast, the density of E2 cells lacking cilia increased with age rising from 40.10 cells/mm^2^ at P10 to 383.9 cells/mm^2^ at P577 (p-value: <0.0001). E2 cells with 4 basal bodies constituted a small subpopulation, whose density remained relatively stable in adult and aged mice. Together, these findings indicate that most E2 cells in the lateral ventricles emerge postnatally, increase in density until P60, and are maintained throughout aging. Interestingly, the composition of the E2 cell population changes substantially with age, with the predominant subtype shifting from uniciliated E2 cells in young animals to E2 cells with no cilia in aged mice.

### Most E2 cells are generated embryonically, but a small number continue to be produced postnatally

The above findings indicate that most E2 cells appear in the lateral ventricles during the first two postnatal months. To determine when E2 cells are generated, we administered a single injection of BrdU to pregnant dams at E13.5, E14.5, E16.5, E18.5, or directly to pups at P4, and analyzed their brains at P30 (n=6). BrdU^+^ E2 cells were identified by their γ-tubulin^+^ donut-like basal bodies and FoxJ1^+^ nuclei (Figure 5A). BrdU^+^ E2 cells were detected following injections at all developmental stages examined. However, more than half of all labeled E2 cells corresponded to the E14.5 injection group (53.7% ± 6.8, cells=654) (Figure 5B). By E16.5, the percentage of double-labeled E2 cells decreased to 19.9% ± 2.5, and it declined further to 2.0% ±2.1 following BrdU administration at P4. These findings suggest that progenitor cells that divide at E14.5 in the ventricular zone give rise to the largest fraction of E2 cells.

**Figure 5.**
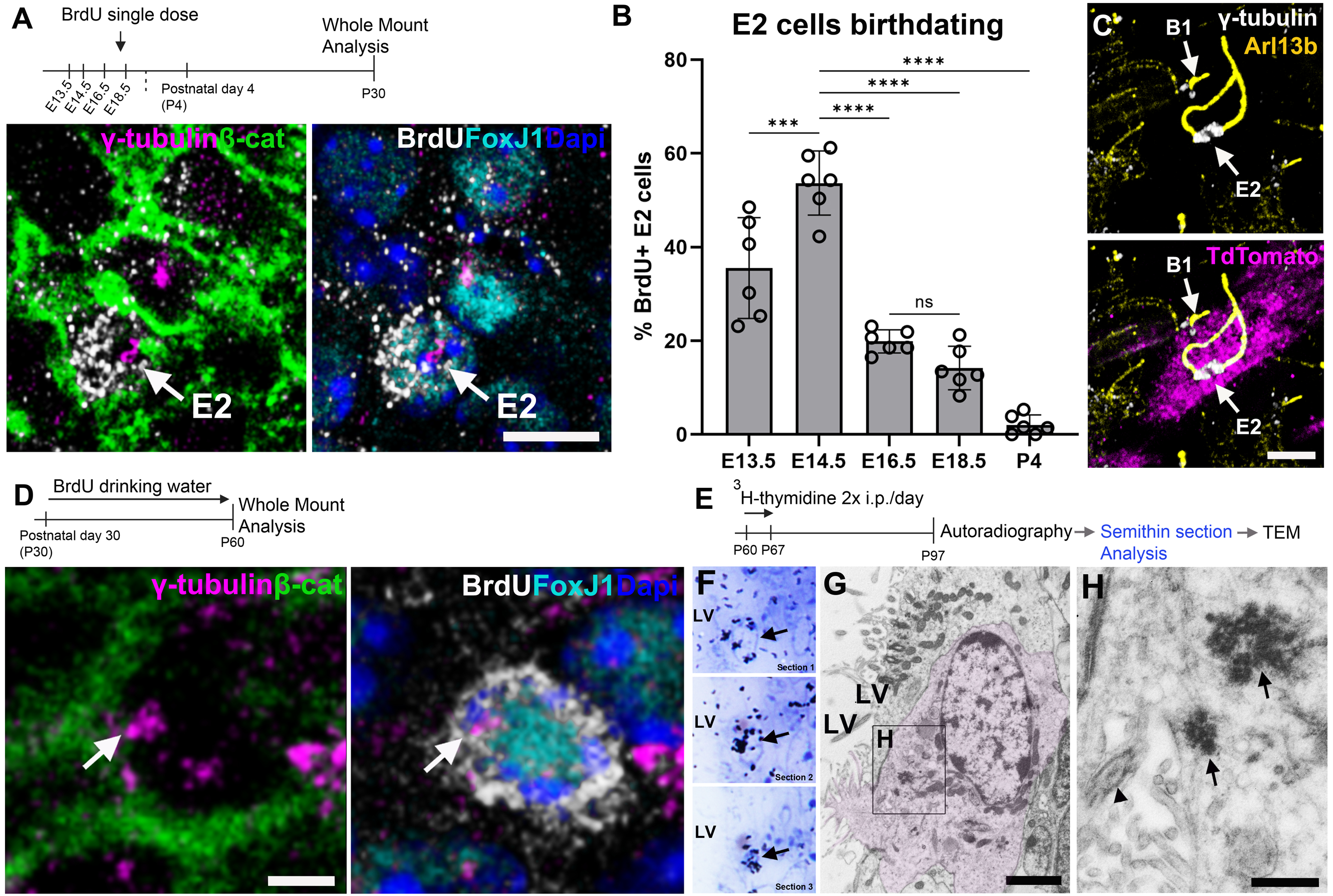
E2 cells are born mostly during embryonic stages (Related to Figure S5). (**A**) Experimental outline for A, B: pregnant females received a single intraperitoneal dose of Brdu at Embryonic stages (E)13.5, 14.5, 16.5 or 18.5 and pups at Postnatal day (P) 4. Whole mounts of the lateral ventricles were analyzed at P30 (n=6 per time point). Confocal images of BrdU^+^ E2 cells identified by γ-tubulin donut-like basal bodies (magenta) and FoxJ1^+^ nuclei (cyan). (**B**) Proportion of E2 cells generated at different developmental stages. More than half of all labeled E2 cells were born at E14.5 (53.7% ± 6.8%; 654 cells). (**C**) Representative confocal image of a P30 lateral ventricle whole mount from a Nestin::CreER;Ai14 mouse that received a low dose of tamoxifen at E14.5, showing a tdTomato+ clone containing a biciliated E2 cell adjacent to a B1 cell. (**D**) Experimental design: P30 mice received BrdU in the drinking water for 1 month. Representative confocal image showing a FoxJ1+/BrdU+ E2 cell. (**E**) Experimental outline: P60 mice received two i.p. injections of 3H-Thymidine (3H-Thy) /day for 7 days, the V-SVZ was analyzed one month after the last injection. (**F**) Autoradiography of 3 consecutive toluidine blue-stained semithin sections (1.5um) of the V-SVZ showing a ^3^H-thymidine^+^ label retaining cell (arrow). (**G-H**) TEM micrographs of the ^3^H-Thy-labeled cell in (F), identified as a biciliated E2 cell showing two enlarged basal bodies (arrows) and a 9+2 cilia (arrowhead). Scale bars: 5μm (A), 3 μm (D), 2 μm (C, G), 500nm (H).

To independently fate-map radial glial cells at E14.5, we administered a single low dose of tamoxifen to Nestin::CreER;Ai14 pregnant females at this age and examined whole-mount preparations from their offspring at P30^51^. By using γ-tubulin and Arl13b to identify E2 cells, we found 14 TdTomato_⁺_ uniciliated and 6 TdTomato_⁺_ biciliated E2 cells (788 E1, 373 B1, and 20 E2 cells; n = 3) (Figure 5C, Figure S5A). At the tamoxifen dose used, labeled cells in the lateral wall of the LV appeared as discrete clusters, likely derived from individual recombined radial glial cells^52^. Tdtomato^+^ E2 cells were frequently located adjacent to Tdtomato+ B1 or E1 cells.

Together, these findings indicate that most E2 cells present at P30 were generated during embryogenesis, with the largest fraction emerging between E13.5 and E14.5. This timing is slightly earlier than E1 cells, whose production peaks between E14 and E16 ^48^. The presence of E1 and B1 cells within clusters containing labeled E2 cells suggest that these cell types can originate from a common progenitor.

A small number of E2 cells were labeled at P30 following BrdU administration at P4, suggesting that E2 cell production continues at a slower rate after birth. To determine if E2 cells are also generated during early adulthood, we administered BrdU in the drinking water to P30 mice for 30 days and analyzed the number of BrdU^+^ E2 cells at P60. Among 236 E2 cells analyzed from four mice (n=4), we found 4 BrdU^+^ E2 cells (1.72 %) (Figure 5D). This is consistent with a small proportion of E2 continuing to be produced after one month of age. To further confirm that some E2 cells were generated in the adult mouse brain, we administered [³H]-thymidine every 12 hours for seven days in P60 animals and analyzed their lateral ventricles 30 days later (Figure 5E). Labeled cells were detected in autoradiograms of 1.5 μm semithin sections under the light microscope (Figure 5F and Figure S5B, C). We selected 19 labeled cells that were in the ventricular zone in the semithin sections. TEM analysis of these 19 cells revealed 2 cells identified as E2 based on their large donut-like basal bodies and 9+2 microtubule organization in their cilia (Figure 5E, F and Figure S5B-G). To determine if E2 cells self-renew, we injected a single dose of Brdu (50 mg/kg weight) in P30 mice and analyzed E2 cells 1 day and 3 days after injection. No labeled Brdu^+^ E2 cells were observed at 1 day or 3 days after injection (d1: 371 E2 cells, n=4; d3: 178 E2 cells, n=4). Together, these data indicate that most E2 cells are generated during embryonic development, and a small fraction continues to be generated in the juvenile and young adult brain.

### E2 cells are Shh-responsive

As shown above, E2-cell cilia express high levels of Arl13b and Inpp5e, two proteins involved in the regulation of Sonic Hedgehog (Shh) signaling ^53,54^. In the canonical Shh pathway, Shh binds to Patched1, relieving its inhibition of Smoothened (Smo) and allowing the translocation and accumulation of Smo into the cilium ^55,56^. We therefore investigated if Shh induced Smo ciliary localization in E2 cells. In basal conditions we found that 21.52 ± 6.91% of E2 showed Smo localization within their cilia (Uni: 28.29 ± 8.08% and Bi: 11.01 ± 6.70%, P30, n=3, 156 cells). To test if Shh stimulation promotes Smo translocation, we cultured whole mounts of the lateral ventricles for 24 hours in a medium containing Shh (Figure 6A). Following treatment, explants were immunostained for β-catenin, Arl13b, and Smo to identify E2 cells and evaluate Smo localization within their cilia. Comparison of vehicle- and Shh-treated explants revealed a marked increase in the proportion of E2 cells showing ciliary Smo localization, from 36.81 ± 22.82% in vehicle-treated explants to 70.67 ± 9.35% in Shh-treated explants (n = 9, 3 independent experiments; vehicle = 419 cells, Shh = 369 cells). This increase was observed in both uniciliated and biciliated E2 cells (Figure 6B, C). Members of the Gli family of transcription factors act downstream of Smo in the canonical Shh signaling pathway^57^. To determine if Smo translocation to E2-cell cilia was associated with downstream pathway activation, we co-immunostained the explants with an antibody against Gli2. We observed an increase in Gli2 expression within E2-cell nuclei following Shh treatment (Figure 6D), suggesting activation of downstream Shh signaling in these cells. These results suggest that E2-cell cilia detect extracellular Shh and transduce canonical Hedgehog signaling.

**Figure 6.**
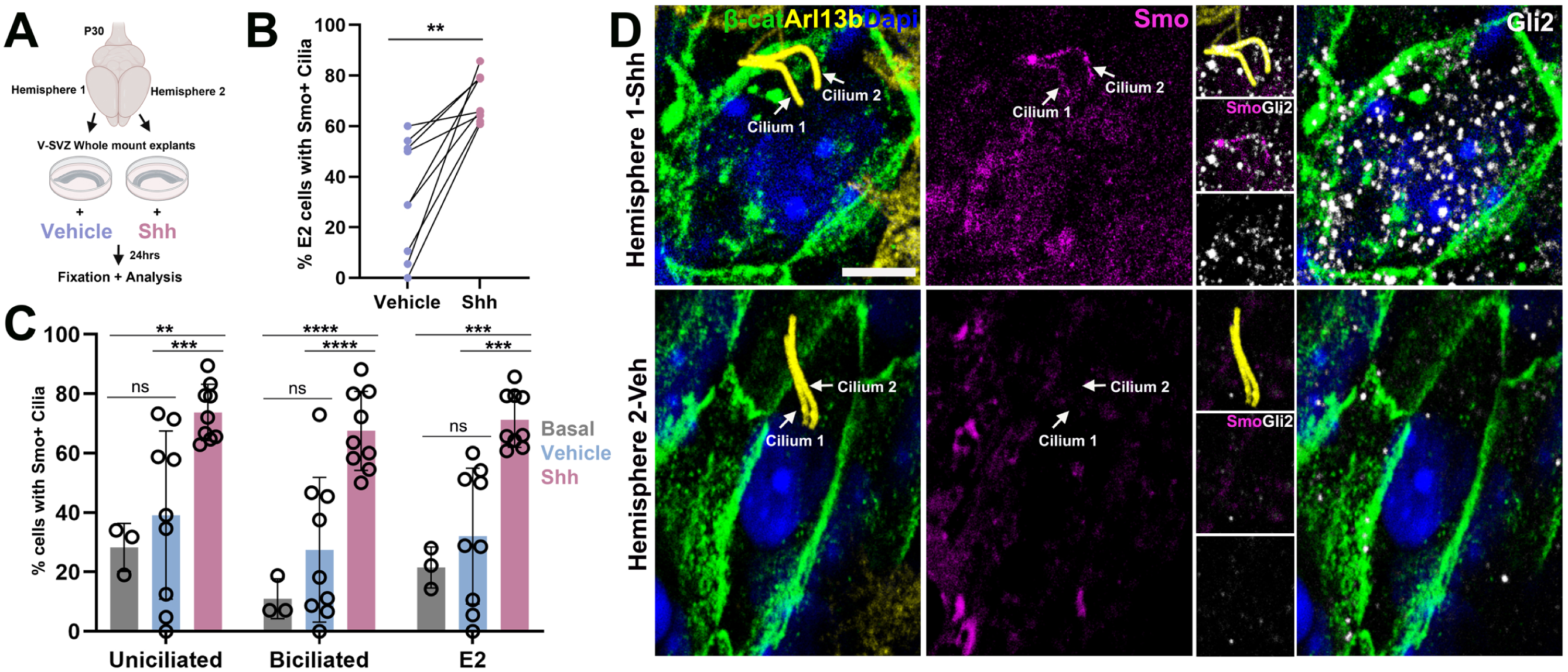
E2 cells respond to Shh signalling. (**A**) Schematic of the experimental design. (**B**) Paired plot showing the percentage of E2 cells with ciliary Smo localization in vehicle- or Shh-treated whole-mount explants. The percentage of E2 cells with ciliary Smo increased from 36.81 ± 22.82% in vehicle-treated explants to 70.67 ± 9.35% following Shh treatment (n = 9; 3 independent experiments; vehicle: 419 cells; Shh: 369 cells). (**C**) Bar plots showing the percentage of E2 cells with ciliary Smo localization under basal conditions (P30, n = 3, 153 cells) and in explants treated with vehicle or Shh (P30, n=9). (**D**) High-magnification confocal images of biciliated E2 cells identified by β- catenin (green) and Arl13b (yellow) from the same brain. Note that Shh-treated E2 cells exhibit Smo (magenta) localization in their cilia and Gli2 (white) accumulation in the soma. Statistical analyses: paired *t* test in (B) and one-way ANOVA with Tukey’s multiple-comparisons test in (C). Statistical significance was defined as ∗∗p < 0.01, ∗∗∗p < 0.001, and ∗∗∗∗p < 0.0001. Scale bar: 5 μm (D).

## Discussion

Here we identified E2 cells as a distinct population of forebrain ependymal cells present in the mouse and human lateral ventricles. E2 cells were characterized by large donut-like basal bodies bearing 2, 1 or no cilia. We show that these unique basal bodies correspond to enlarged subdistal appendages. Interestingly, E2 cilia exhibited a unique pattern of linear motility that differed from that of multiciliated E1 cells. E2 cells were distributed throughout the lateral ventricular wall, with higher densities in regions of high CSF flow and neurogenesis ^11,15,58^. E2 cells were generated predominantly during embryonic development and completed their differentiation postnatally, although a small fraction continued to be produced shortly after birth and, rarely, in adulthood. Following their postnatal differentiation, E2 cell numbers increased during young adulthood and then remained relatively stable, whereas the number of cilia per cell declined with age. Their distinctive ciliary architecture, motility pattern, molecular profile and anatomical localization suggested a sensory role. Consistently, E2 cells responded to Shh signaling by increasing Smo accumulation in their cilia. This novel population of ependymal cells is likely to have important functions on the regulation of CSF signaling.

Our results indicate that E2 cell cilia combine structural and molecular properties of both motile and primary cilia. The ciliary shaft of E2 cells contains a 9+2 axoneme with radial spokes and expresses Spef2, as well as Dnah9 and Ccdc33 at the ciliary tip, all hallmarks of motile cilia ^23,24,59^. Consistent with these features, live imaging demonstrated that E2 cilia were motile. However, E2 cilia also expressed high levels of Arl13b and Inpp5e, two key proteins associated with primary cilia and Shh signaling^60–62^. Moreover, the mother centriole of uniciliated E2 cells displayed 9 radially organized sDAPs and remained linked to the daughter centriole through intercentriolar fibers, a basal body organization characteristic of primary cilia. In contrast, the basal bodies of classical motile cilia, such as those of E1 cells, typically contain a single basal foot (a modified sDAP) and are not tethered to daughter centrioles^29,35,63^. The distinctive organization of the E2 basal body further supports the notion that these cells harbor specialized cilia. Their basal bodies are approximately 45-fold larger than those of E1 cells, and we show here that this remarkable morphology results from exceptionally large and elaborated sDAPs. To our knowledge, such enlarged sDAPs have not been described in other tissues^25,26,64^. sDAPs mediate microtubule anchoring and participate in vesicular trafficking involved in ciliary signaling ^64–66^. Consistent with these functions, we observed a dense radial array of microtubules emerging from the sDAPs of E2 cells, frequently associated with small vesicles and multivesicular bodies. Together, these findings suggest that E2 cells may detect and transduce information about the molecular composition of the CSF.

Live imaging showed that E2 cilia were motile, but this motility was very different from that described in E1 cells^36^. The proximal region of E2 cilia moved linearly back and forth, hinged at the basal body. The distal portion of E2 cilia moved more freely in a whiplash manner, influenced by both the movement of the proximal region and the flow of CSF. Unlike E1 cells, which draw the distal segment of their cilia close to the apical surface during the recovery stroke to minimize backward CSF flow, E2-cell cilia remained extended throughout the beat cycle. The one or two cilia in E2 cells, compared to the ∼50 cilia of E1 cells, would have only a minimal effect on the overall flow of the CSF. Rather than propelling the surrounding fluid, the motility in E2 cilia is likely exploratory. This is consistent with E2 cells having a sensory function (chemical, mechanical or a combination of both). The back-and-forth movement could help sample a larger volume of CSF compared to a stationary cilium. The observation that biciliated E2 cells show asynchronous beating, together with the distinct basal body architectures of the mother centriole (full SDAs ring) and daughter centriole (horseshoe configuration), also raises the intriguing possibility that the two cilia are structurally and functionally specialized. Biciliated E2 cells could differentially detect and integrate distinct mechanical or chemical cues from the CSF, thereby expanding the sensory repertoire in a single E2 cell.

BrdU birth-dating experiments revealed that most E2 cells are generated during embryonic development, with peak production around E14.5. A smaller fraction continued to be generated during late embryonic and postnatal stages with a very small number still produced in adulthood. This developmental window overlaps with the period during which both E1 cells and B1 cells (NSCs) are generated^48,51,67^. Consistent with this timing, our lineage-tracing analysis in Nestin::CreER;Ai14 mice indicate that E2 cells also arose from the same pool of radial glial progenitors. This finding agrees with previous lineage-tracing studies showing that individual radial glial cells can generate both E1 and B1 cells^52,68^. Our results expand the repertoire of radial glia-derived ependymal cells to include E2 cells^69^. Although E2 cells share a common developmental origin with E1 cells, their maturation appears to follow a slower timeline. Whereas most E1 complete their maturation before P14 in mice^48,49^, a subset of E2 cells continues to be generated and mature until at least P60. In adult mice, we did not detect any labeled E2 cells 1 or 3 days following BrdU administration, whereas we detected a small number of labeled E2 cells 1 month after Brdu or 3H-Thymidine labeling. These results suggest that most E2 cells become postmitotic following their embryonic generation, like E1 cells^48,70^, and that continued increase in mature E2 cells reflects delayed differentiation, with only a small contribution from a source other than the division of postnatal E2 cells. It is likely that the few E2 cells produced in adulthood derive from proliferating local NSCs (V-SVZ B1 cells), although this remains to be demonstrated.

Mapping E2 cells along the lateral ventricles revealed spatial heterogeneity. All four E2 cell populations were enriched in the anteroventral and posterodorsal regions. These areas are known sites of high neurogenic activity^45^, and respond to physiological states such as pregnancy^58^, as well as hunger and satiety signals^71^. Intriguingly, both regions are associated with restricted territories of Shh signaling. The anteroventral region shows active Shh signaling in adulthood ^72^, while the posterodorsal corridor exhibits transient Shh signaling during the first two postnatal weeks ^73^, coinciding with the onset of E2 maturation. Consistent with a potential association between E2 cells and Shh signaling, we found that E2-cell cilia express substantially higher levels of Arl13b and Inpp5e than E1 cells, reaching levels comparable to those observed in the primary cilia of B1 NSCs. Both Arl13b and Inpp5e have been implicated in the regulation of Shh signaling during development^53,54^. Supporting a functional role for E2-cell cilia in Shh signalling, a small subset of E2 cells exhibited ciliary Smo localization under basal conditions, and this proportion increased significantly following Shh stimulation. Further studies will be required to determine how Shh signaling influences E2 cell function, as well as the source and regulatory role of Shh in the adult CSF.

Here, we show that the walls of the lateral ventricles, in both mice and humans, contain a population of ependymal cells with unique cilia and basal bodies. Their distribution, ciliary structure and motility suggest that these cells have a sensory function. Interestingly, E2-like cells, with one or two cilia and large basal bodies, have also been detected in the walls of the third and fourth ventricles in mice and humans^17,74^ and in the central canal of the spinal cord of mice, macaques, and humans^18,19^. It remains unknown whether E2-like cells throughout the ventricular system constitute a conserved CSF-sensing cell type or are regionally specialized and tuned to the molecular composition of the different ventricular compartments, while sharing a similar basal body-cilium architecture. Comparative molecular analyses of E2 cells across the ventricular system will help define the function of E cells and determine how E2 cells differ along the neuroaxis. Further defining the molecular identity of E2 cells will help determine how they contribute to CSF homeostasis and disease. Interestingly, E2 cells may also be relevant to understand commonly diagnosed brain tumors. Intriguingly, Grade II ependymomas, arising in the posterior fossa and fourth ventricle, have been reported to contain cells with enlarged basal bodies and zero, two, or three 9+2 cilia that resemble the E2 cells^75^. Together, our findings identify E2 cells as an evolutionary conserved sensory ependymal cell subtype and provide a foundation for investigating their roles in human brain’s ventricular biology and disease.

### Limitations of the study

E2 cells represent a small fraction (<5%) of all ependymal cells^45^. Although they can be readily identified *in situ* by their distinctive morphology, E2 cells cannot yet be isolated or enriched for molecular and functional analyses. Their low abundance and the lack of specific molecular markers have limited our ability to define their unique molecular profile. Additional analyses of our previously published unbiased single-cell RNA sequencing datasets, together with extensive validation in tissue (unpublished data), did not identify a distinct E2 cell cluster. These findings suggest that the rarity of E2 cells currently limits their confident molecular characterization, and that larger single-cell datasets or targeted enrichment strategies will likely be required to resolve their molecular identity. To distinguish the unique motility pattern of E2 cilia we used live imaging of acute whole mount explants maintained for 1-5 hours at room temperature. The speed of motility could differ *in vivo* within the biological context of a closed ventricle and at 37°C. We also cannot exclude effects of neighboring cells within our explants to the observed response to Shh in E2 cells. Our observations using postmortem samples of the human brain lateral ventricular wall only include small fragments of this wall. While these samples reveal E2 cells in the human forebrain ventricular system, we do not know their distribution or abundance; systematic sampling of the walls of the lateral ventricles in humans would be required to determine their overall distribution and abundance.

## Acknowledgments

We thank Professor Semil Chosky and his lab members at UCSF for useful discussions and sharing the Dnah9 antibody. Work in the Alvarez-Buylla laboratory is supported by the Program for Breakthrough Biomedical Research, which is partially funded by the Sandler Foundation, NIH grants R01 NS113910, R01 NS028478, R35 NS143000, and a generous gift from the John G. Bowes Research Fund. FDH and SAR were supported by NIH grant 1K99NS121273. CRC was supported by NIH grant K08 NS126573. We thank Nick Kilner-Pontone and Kenneth Xavier Probst for preparing the illustrations included in Figure 2. We dedicate this work to José Manuel García-Verdugo, as this was one of the last projects to which he contributed his extraordinary expertise in TEM.

## Supplemental Information

Document S1. Figures S1-S5

Table S1. Human samples

Movie S1 and S2

## Methods

### Mice

Mice were housed on a 12 hr day-night cycle with free access to water and food in a specific pathogen-free facility in social cages (up to 5 mice/cage) and treated according to the guidelines from the UCSF Institutional Animal Care and Use Committee (IACUC) and NIH. All mice used in this study were healthy and immuno-competent and did not undergo previous procedures unrelated to the experiment. CD1-elite mice, Fvb/N, C57BL/6 J, Arl13b-mCherry; Centrin-GFP, FoxJ1-CreERt2, and Ai14 mouse lines (see Resources table) were used. Biological and technical replicates for each experiment are described in the relevant subsections below.

### Human Samples

Autopsy consent and all protocols were approved by the Human Gamete, Embryo, and Stem Cell Research Committee and the IRB at UCSF.

### Tissue preparation

Mice were deeply anesthetized with 2.5% Avertin delivered IP. For wholemount preparations, brains were removed and the lateral walls were immediately dissected out as previously described ^11^. Wholemounts were fixed in 4% paraformaldehyde (PFA) at 4°C overnight and then stored in phosphate buffered saline (PBS) containing 0.1% sodium azide at 4°C until immunohistochemical processing.

### BrdU, [3H] Thymidine and Tamoxifen administration

#### BrdU drinking water

To detect label-retaining cells, BrdU (Sigma, cat. B5002) was dissolved in water at 1 mg/ml and administered in the drinking water for 30 days.

#### BrdU injections

BrdU was dissolved in sterile medical grade saline for a final concentration of 5mg/mL. Mice were administered an IP injection with 50mg/kg of BrdU.

#### [3H]-thymidine injections

To identify E2 label retaining cells by TEM, we administered two intraperitoneal injections of 3H-thymidine (1.67 μl/g body weight, specific activity 5Ci/mmol; Perkin Elmer, USA) 12h apart for seven days (P60, n=3 mice). Mice were perfused with 2% PFA-2.5% glutaraldehyde four weeks after the last injection and samples were processed for TEM analysis.

For embryonic Tmx administration, two- to three-month-old, timed-pregnant *Nestin::CreER;Ai14* females received a single dose of Tmx, 1.67mg/kg via oral gavage.

#### For adult Tmx administration

P30 *FoxJ1-CreERt2; Ai14* mice received 100 mg/kg via oral gavage in corn oil.

### Whole-Mount explants

Whole mounts of the V-SVZ of the P30 CD1 mice were performed under dissection microscope and L15 Medium. Per each mouse, one whole mount was placed on a 48 well plate in DMEM and Fetal bovine Serum, FBS (Hydroclone) 10% and 1% Penicillin/Streptavidine with 200nm Recombinant Mouse Sonic Hedgehog/Shh (C25II) N-Terminus (464-SH-025) in BSA 1% (right hemisphere) or BSA 1% (vehicle, left hemisphere) during 24 hours into a 37 C, 5% CO2 and 8% O2 incubator. After 24 hours samples were rinsed with sterile PBS and fixed with PFA 4%.

### Immunohistochemistry

Wholemounts were incubated in primary and secondary antibodies in TNB Blocking Buffer (0.1 M Tris-HCl pH 7.5, 0.15 M NaCl, 0.5% blocking reagent) Akoya Biosciences overnight at 4°C. For samples processed for BrdU unmasking (Citrate solution, 20min), incubations with primary and secondary antibodies were performed first, followed by fixation in 4% PFA at room temperature for 10 minutes. Tissue was then incubated in 2N HCl at 37°C for 30 minutes, followed by treatment with 0.1M boric acid (pH8.5) for 20 minutes at RT prior to incubation with anti-BrdU antibodies and corresponding secondary antibodies for 24 hrs at 4°C each. Primary antibodies are listed on the methods table. Secondary antibodies were conjugated to Donkey AlexaFluor dyes (Invitrogen/Molecular Probes).

### Transmission electron microscopy

For TEM, mice were fixed with 2% PFA-2.5% glutaraldehyde. Brains were rinsed in 0.1 M phosphate buffer (PB) and cut into 200μm sections. Sections were post-fixed in 2% osmium tetroxide, dehydrated, and embedded in Durcupan resin (Fluka; Sigma-Aldrich). Semithin sections (1.5μm) were cut with a diamond knife and stained with 1% toluidine blue for light microscopy. Ultrathin sections (70–80 nm) were cut, stained with lead citrate, and examined under an FEI Tecnai G2 Spirit transmission electron microscope (FEI Europe) using a digital camera (Xarosa (20 Megapixel resolution), Radius EMSIS GmbH, Münster, Germany). Immunogold staining and TEM processing were performed as previously described ^76^.

### TEM serial reconstructions

For uni- and bi-ciliated E2 cell serial reconstructions, all serial ultrathin sections (∼100 sections of 70 nm thickness each) from either an en face or coronal view of the V-SVZ were imaged. As a criterion to identify E2 cells, all reconstructed cells exhibited one or two cilia and at least one large, electron-dense basal body. Serial images of basal bodies were sequentially ordered in Photoshop (Adobe), and the paint bucket tool was used to pseudo-color (magenta) the pixels corresponding to the highest electron density. For 3D reconstruction models, serial TEM micrographs were stacked and aligned using FIJI TrakEM2 software. TEM stacks were rendered with the surface tool in Imaris software (v10).

### Autoradiography

Brains injected with 3H-Thy were processed for TEM as described above. Subsequently, V-SVZ semithin sections were dipped in autoradiography emulsion (Carestream Autoradiography Emulsion, Type NTB), dried in the dark, and stored at 4°C for 4 weeks. Autoradiography was developed using standard methods and counterstained with 1% toluidine blue. Close to the LV 3H-Thy-labeled nuclei were identified in semithin sections. For a cell to be considered labeled, six or more silver grains needed to be present over the nucleus, and the labeled nucleus needed to be observed in at least three consecutive serial sections. All consecutive sections showing labeled cells were selected under a light microscope (Eclipse; Nikon), re-embedded, and ultrathin-sectioned (70nm) for TEM analysis (Tecnai Spirit G2, FEI, Oregon). Anatomical references were used to identify labeled cells under EM. E2 cells were identified by the presence of large electron-dense basal bodies associated to one or two cilia with 9+2 axonemal organization.

### E2 cell subtype map

A wholemount (P60, FVB/NJ female mouse, n=1) was dissected, fixed, and stained for γ-tubulin, β-catenin, Arl13b. Confocal Z-stacks of 211 V-SVZ positions were obtained with a 63x oil objective. The beginning plane for z-stacks was set on the ependymal cilia tips (identified by Arl13b) to ensure that all apical cells were included in the analysis. The end plane for each stack was set to include the whole thickness of the SVZ. We considered that we reached the striatum when the DAPI cells were no longer aggregated in cell clusters (Z size was variable between regions, Z step size=0.5). Images were stitched and analyzed with Imaris software. Epithelial cells in the lateral wall were identified based on their apical surface, delineated by β-Catenin staining, and the number and size of their basal bodies marked by γ-tubulin ^11^. Subtypes of E2 cells were annotated using the Imaris Spots tool, based on the number of large ring-like basal bodies and the number of cilia. To visualize the spatial distribution of E2 cells subtypes along the lateral wall, spot position data was exported to R. Spatial maps of E2 cells subtypes were generated using the ggplot2 package.

### Quantifications

#### Cilia length and Arl13b fluorescence intensity

Wholemounts from CD1 mice (P33, 2 females and 1 male) were dissected, fixed, and immunostained for β-catenin, γ-tubulin and Arl13b. Confocal Z-stacks of 6-9 positions/region (AV and PD)/mouse were obtained using a 63x oil objective. E2 cells were identified by their long Arl13b bright cilia. Arl13b stained cilia were then reconstructed using the Filaments function on the Imaris 10.1.0 software following the entire length of the cilia through the Z-stack. Measurements of “intensity mean” and length of the reconstructed filament were recorded for each cilium.

#### Planar polarity quantifications

Confocal images of E2 cells were imported and analyzed with FIJI. ß-Catenin-stained surface area of the E2 cells was traced with the freehand tool and measured its center of mass coordinates. The surface area of basal bodies and their center of mass coordinates were measured likewise. To measure the displacement of the basal bodies, the radius from the center of the cell to the center of mass of the basal bodies was traced along with the radius from the center of the cell to the surface in the same direction using the line tool. Measurements of orientation of E2 basal bodies with respect to CSF flow were done by tracing a vector following the orientation of E1 basal bodies as 0 degrees.

#### E2 cell subtypes over time

Whole mounts from FVB/NJ mice (P10, P21, P28, P60, P90, P120, P180 and P365 (n= 3/age/sex) and P545 (3 females, 1 male)) were dissected, fixed, and immunostained for β-catenin, γ-tubulin and Arl13b. Confocal Z-stacks of 9 positions/region (antero-ventral and posterior-dorsal)/mouse were obtained using a 63x oil objective. E2 cells with 1, 2 or no cilia and 4 basal bodies were identified by their donut-shaped γ-tubulin basal bodies.

#### Adult BrdU drinking water

Wholemounts from CD1 mice (P60, 4 females) were dissected, fixed, and immunostained for BrdU, γ-tubulin, FoxJ1, β-catenin, and DAPI. Confocal Z-stacks of 4 positions/region (AV and PD)/mouse were obtained using a 63x oil objective. E2 cells were identified by their donut-shaped basal bodies and FoxJ1^+^ nuclei, then marked as either BrdU+ or BrdU-.

#### Embryonic BrdU Injections

Wholemounts from offspring of BrdU injected dams were dissected and fixed at P30 (n=3/sex/time of injection). Wholemounts were immunostained for BrdU, γ-tubulin, FoxJ1, -catenin, and DAPI. Confocal Z-stacks of 4 positions/region (AV and PD)/mouse were obtained using a 63x oil objective. E2 cells were identified by their donut-shaped basal bodies and FoxJ1+ nuclei, then marked as either BrdU+ or BrdU-.

##### Smo expression in whole-mount explants

Samples were de-identified prior to imaging and analyzed in a blinded manner (3 independent experiments, 3 mice/experiment (n=9), P30 CD1). De-identified specimens were imaged using confocal microscopy. The surface of each whole mount was scanned, and E2 cells were identified by their Arl13b-bright cilia. Z-stacks extending from the ciliary tip to the E2 cell soma were acquired for each fluorescence channel with 0.5 µm step intervals. Each channel was imaged sequentially using restricted fluorescence wavelength ranges to minimize spectral crosstalk. Images were manually quantified to determine Smo colocalization with Arl13b along the length of E2 cell cilia. For each sample, the percentage of E2 cells exhibiting ciliary Smo localization was calculated relative to the total number of E2 cilia analyzed. Following completion of quantification, samples were de-identified to their experimental groups for statistical analysis.

### Motility Analysis

Whole mounts of the ventricular lateral wall from P30 Arl13b-mCherry; Centrin-GFP mice were dissected and placed in oxygenated artificial cerebrospinal fluid (aCSF) containing 110 mM choline chloride, 2.5 mM KCl, 7 mM MgCl2, 0.5 mM CaCl2, 1.3 mM NaH2PO4, 25 mM NaHCO3, 10 mM D-(+)-glucose and 1× penicillin–streptomycin. Wholemounts were secured to a cell culture dish with oxygenated aCSF for stable recording for a maximum of 5 hours. E2 cells were identified using centrin and their long Arl13b bright cilia in the anteroventral and posterodorsal regions of the ventricular lateral wall. Recordings of ciliary movement were taken using a 25x dipping objective on the Nikon A1R, an upright laser scanning confocal microscope. Videos were captured with 6x zoom, frame time 130.0 ms, 7.7 frames/second, 1.01 µm/px, and 512x512 pixels. At least 3 videos/E2 cell were captured at different Z-planes across the length of the cilia to record movement at the proximal segment, midpoint, and the very tip of the cilia. Videos of E2 ciliary movement were then analyzed on the Imaris 10.1.0 software using the “Filament” and “Spot” tools. For videos showing the proximal segment, a spot was placed about 3µm from the base of the cilium in each frame (115-154 frames/video, 7.7 fps in 15-20 sec videos). For videos showing the tip of the cilium, a spot was placed at the end of the cilium in each frame. Positional data of the location of the proximal segment and the tip in relation to the base of the cilium for the duration of the video was exported to R. Plots showing position and number of beats/second of cilia were generated using ggplot2.

### RNAscope assay

For *en face* sections, CD1 Mouse brains (P30) were serially sectioned using a Leica cryostat (12 μm-thick sections in Superfrost Plus slides) (n=2-4 mice/probe). Sections were incubated 10 min with 4% PFA and washed 3x10 min with phosphate-buffered saline (PBS) to remove OCT. Slides were incubated with ACD hydrogen peroxide for 10 min, treated in 1x target retrieval buffer (ACD) for 5 min (at 96–100 °C), and rinsed in water and 100% ethanol. Samples were air dried at 60°C for 15 min and kept at room temperature overnight. The day after, samples were treated with Protease Plus for 10 min at 40°C in the RNAscope oven. Hybridization of probes and amplification solutions was performed according to the manufacturer’s instructions. Amplification and detection steps were performed using the RNAscope Multiplex kit. RNAscope assay was directly followed by antibody staining for anti-ß- Catenin and mouse anti-γ-tubulin. Samples were blocked with TNB solution (0.1 M Tris–HCl, pH 7.5, 0.15 M NaCl, 0.5% PerkinElmer TSA blocking reagent) for 30 min and incubated in primary antibodies overnight. Samples were washed with PBS-Tx0.1% and incubated with secondary antibodies Donkey anti-Chicken 488 (Jackson ImmunoResearch, 1:500) and Donkey anti-mouse 647 (Jackson ImmunoResearch, 1:500) in TNB buffer for 1.5 hrs. Washed 3x5 min and incubated with DAPI 10 min. Sections were mounted with Prolong Glass Antifade Mountant (Invitrogen, P36980).

### Statistics

All results shown in the graphs are expressed as mean ± SD. Statistical significance was defined as ∗ p ≤ 0.05, ∗∗ p < 0.01, ∗∗∗ p < 0.001, ∗∗∗∗ p < 0.0001. One-way ANOVA with Tukey’s multiple comparisons (Prism GraphPad software) was used to determine the statistical significance of multiple comparisons; Student’s t-test (Excel) was used for pairwise comparisons between two groups, and paired test t to compare Shh treated and Shh non-treated same groups.

## Resources Table

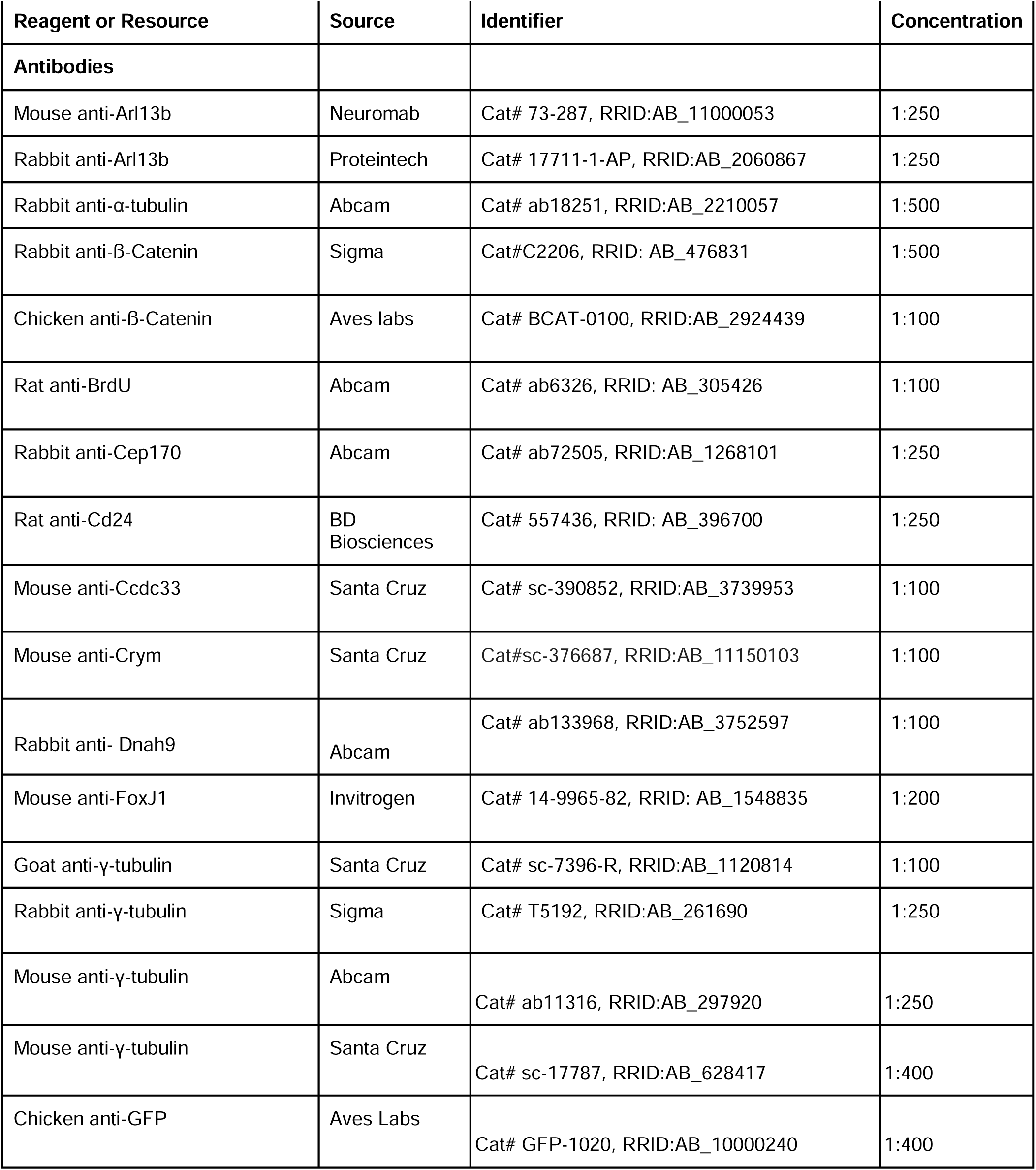
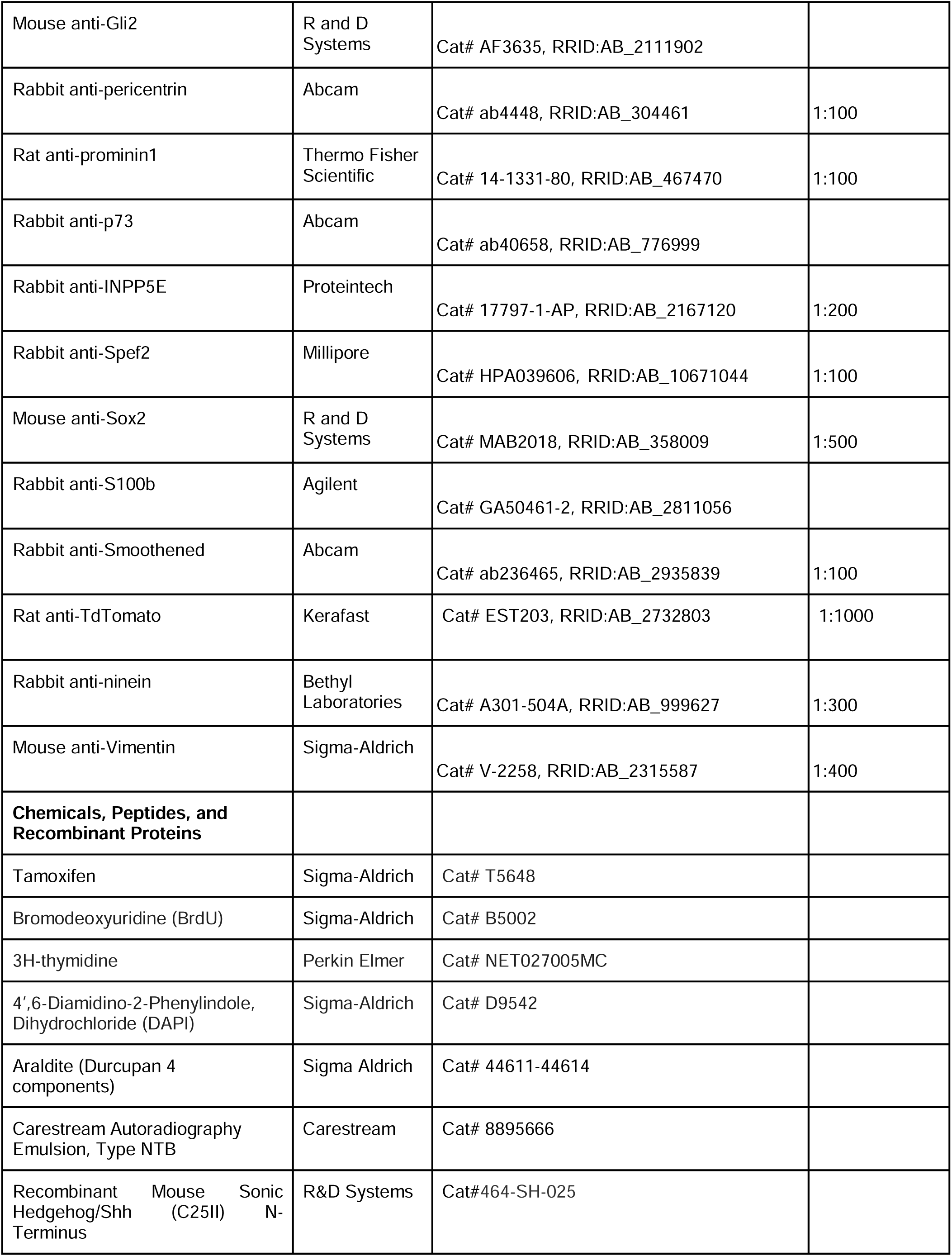
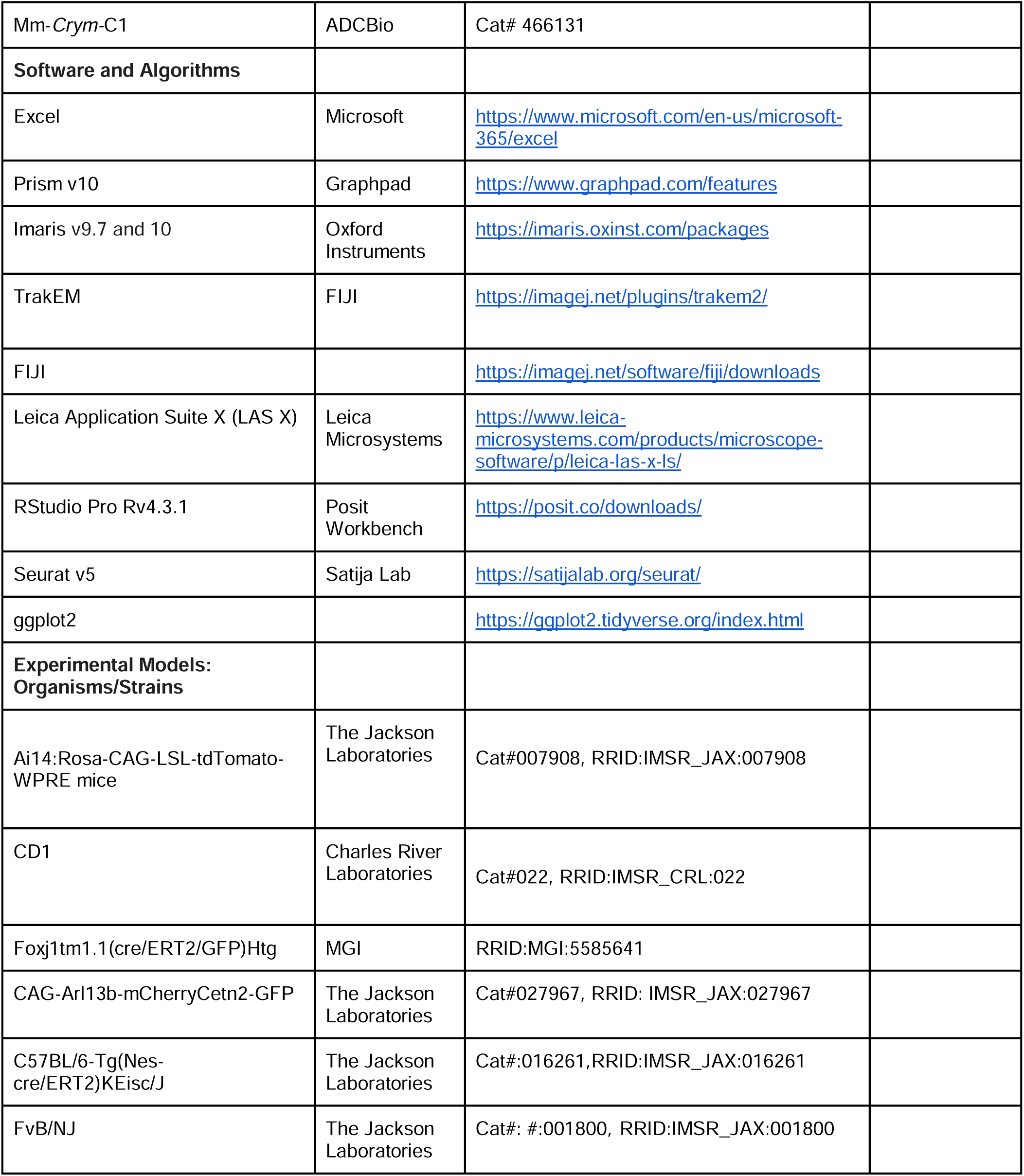

