## Supplemental Figures for "Specialized Shh-sensing cell with unique cilia and basal body in the forebrain ventricular epithelium"

#### Supplementary Figure 1

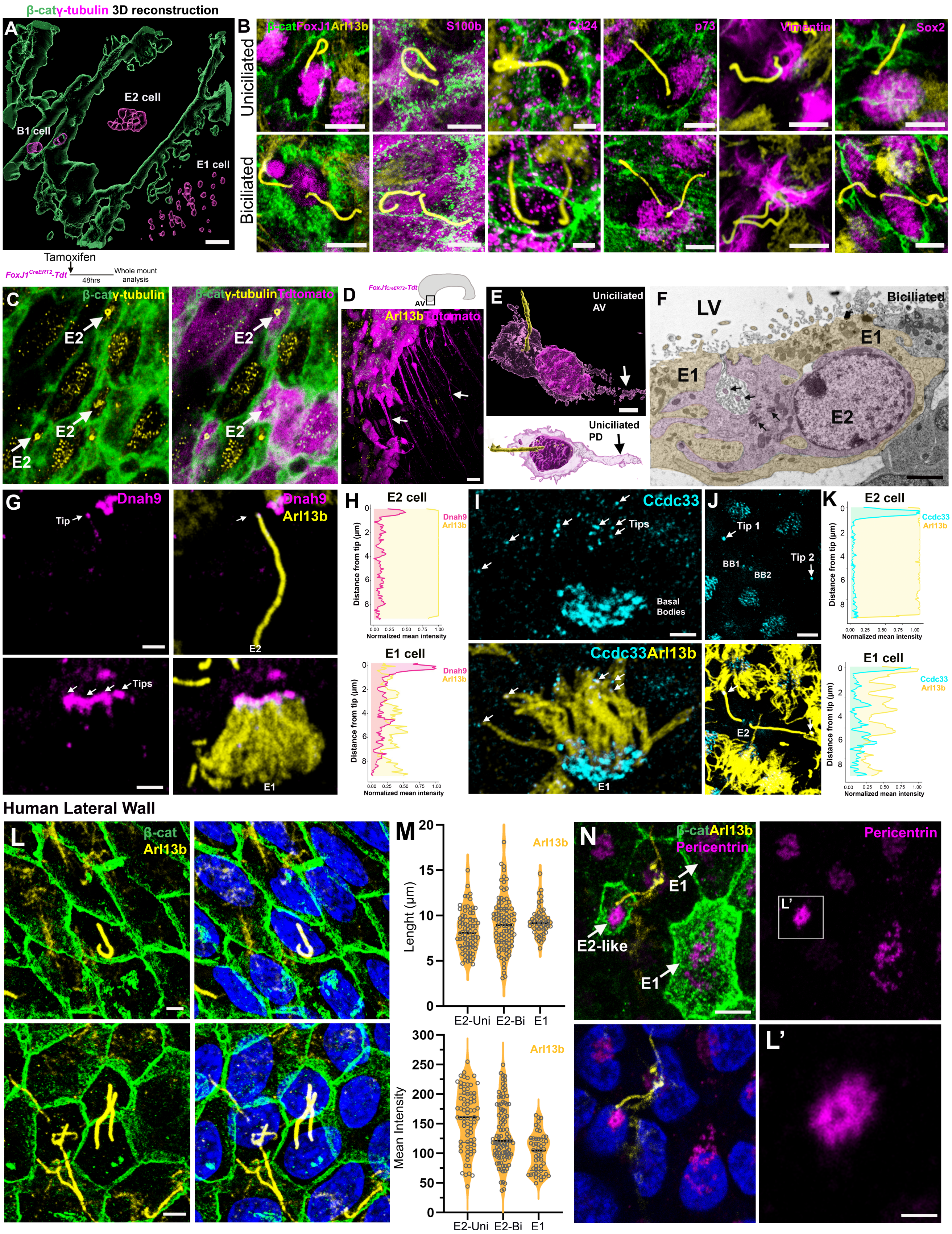


**Figure S1. Uni- and Biciliated E2 cells molecular expression and morphology.** (**A**) Imaris 3D reconstruction of γ-tubulin (magenta) and β-catenin (green) expression showing the basal bodies organization and size of E1, B1 and E2 cells. **B**) High magnification confocal images of uni- and biciliated cells identified by Arl13b (yellow) and Bcat (green) marker expression of FoxJ1, S100b, Cd24, p73, vimentin and Sox2. **C–D)** Confocal images of whole-mount preparations from Foxj1-CreERT2;Ai14 mice after tamoxifen induction showing tdTomato+ (magenta) E2 cells, identified by their characteristic donut-like γ-tubulin+ basal bodies. **(D)**Maximum-intensity projection of the anteroventral lateral ventricular wall showing Foxj1+ ependymal cells with long basal processes. **E)** Imaris 3D reconstructions of Foxj1+ E2 cells illustrating lateral cytoplasmic expansions. **F)** Transmission electron micrograph of an E2 cell (pseudocolored magenta) showing large electron-dense basal bodies (arrows) and two 9+2 cilia (arrowheads). Note the highly interdigitated E2 cell cytoplasm and the nucleus embedded between neighboring E1 cells (pseudocolored brown). **G–H)** Confocal images of Dnah9 localization at the tip of E2 and E1 cells cilia. **(H)** Normalized Dnah9 fluorescence intensity measured along E2 and E1 cilia, using Arl13b to define ciliary length. **I–K)** Confocal images of Ccdc33 localization in E2 and E1 cells cilia. **(K)** Normalized Ccdc33 fluorescence intensity measured along E2 and E1 cilia, using Arl13b to define ciliary length. **L)** Confocal images of human lateral ventricular wall whole mounts stained for Arl13b and β-catenin showing E2-like cells bearing one or two brightly Arl13b-positive cilia. **M)** Quantifications of the length and mean intensity of E2-like cells' cilia. **N)** Confocal image of a whole-mount preparation from the human lateral ventricular wall. An E2-like cell was identified by its donut-like pericentrin-positive basal body and smaller apical surface compared with neighboring E1 cells. Scale bars: 1 μm (A), 2 μm (B, F, G, I, L), 3μm (E), 8 μm (J, N), 15 μm (D).

#### Supplementary Figure 2

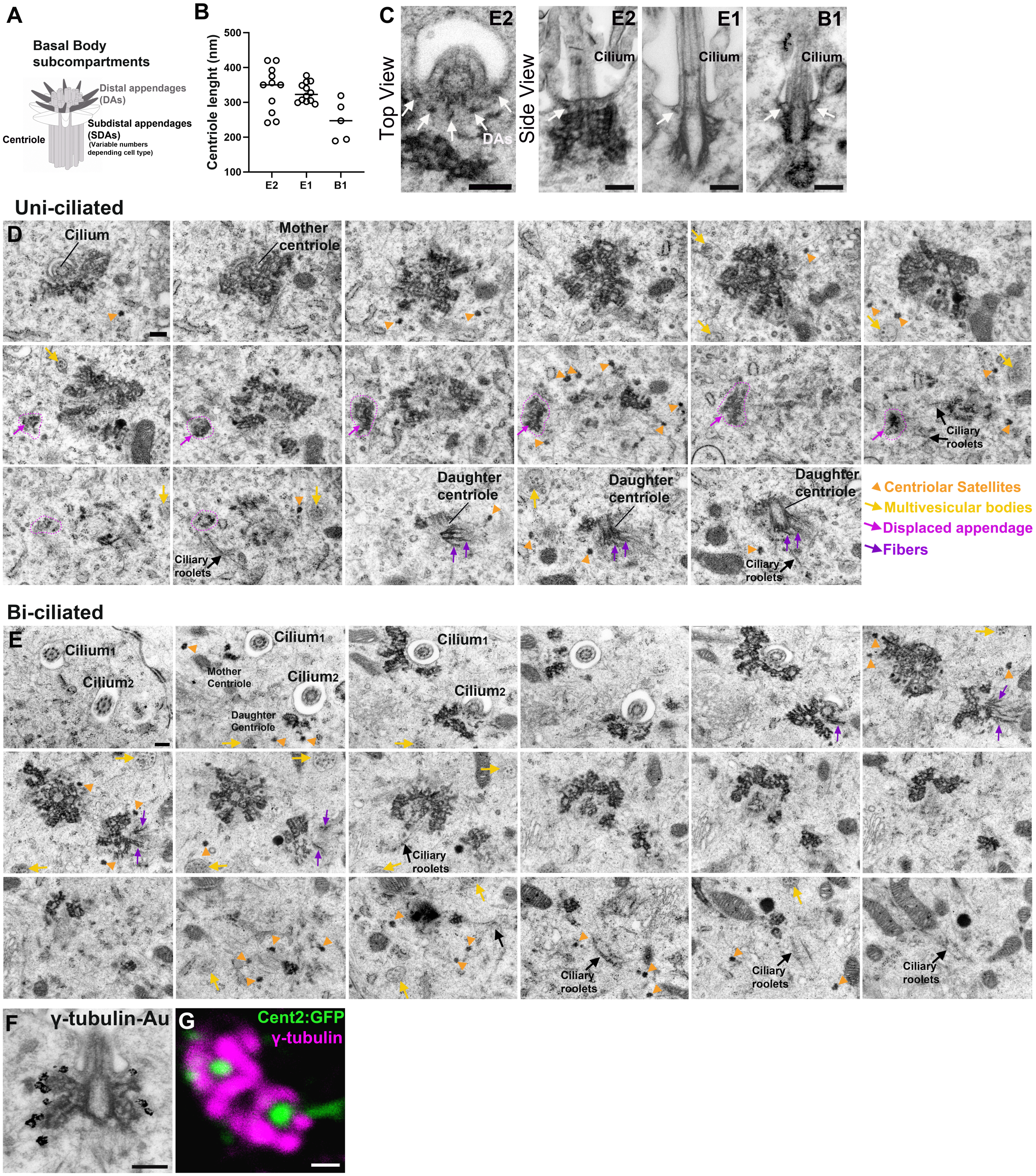


**Figure S2. Basal body reconstructions in Uni- and Biciliated cells.** Related to Figure 2. (**A**) Diagram of basal body subcompartments. Adapted from Ma et al., 2023. (**B**) Quantifications of the centriole length of E1, E2 and B1 cells under TEM. **C**) TEM Micrographs of basal bodies in E2, E1 and B1 cells show distal appendages (DAs) associated with the ciliary membrane (Arrows). (**D, E**) TEM en face serial sections of uni- and bi-ciliated E2 cells in the V-SVZ. E2 cells showed large, electron-dense subdistal appendages (sDAPs). (**D**) Uni-ciliated cells showed one cilium associated with the mother centriole exhibiting 9 large, electron-dense sDAPs. This mother centriole was connected to a daughter centriole with one subdistal appendage and associated fibers.

Electrondense structures with similar organization to the subdistal appendage were close, but not associated with the centriolar wall (magenta arrow and dashed line). Centriolar satellites (orange arrowhead) and multivesicular bodies (yellow arrows) were frequently associated with E2 cell basal bodies. (**E**) In bi-ciliated cells two cilia are present. One originated from a centriole with 9 large, electron-dense sDAPs, identified as the mother centriole. The second originated from a centriole with fewer sDAPs and  fibers, identified as the daughter centriole. **F)** Immunogold staining of gammatubulin co-localizing with an E2 cell basal body SDA. G) High magnification confocal image of E2 cells basal bodies labeled by γ-tubulin (magenta) and GFP (green). Note that the donut like and horse shoe organization. Scale bars: 250 nm (C, D), 200 nm (D) and 500 nm (F).

### Supplementary Figure 3

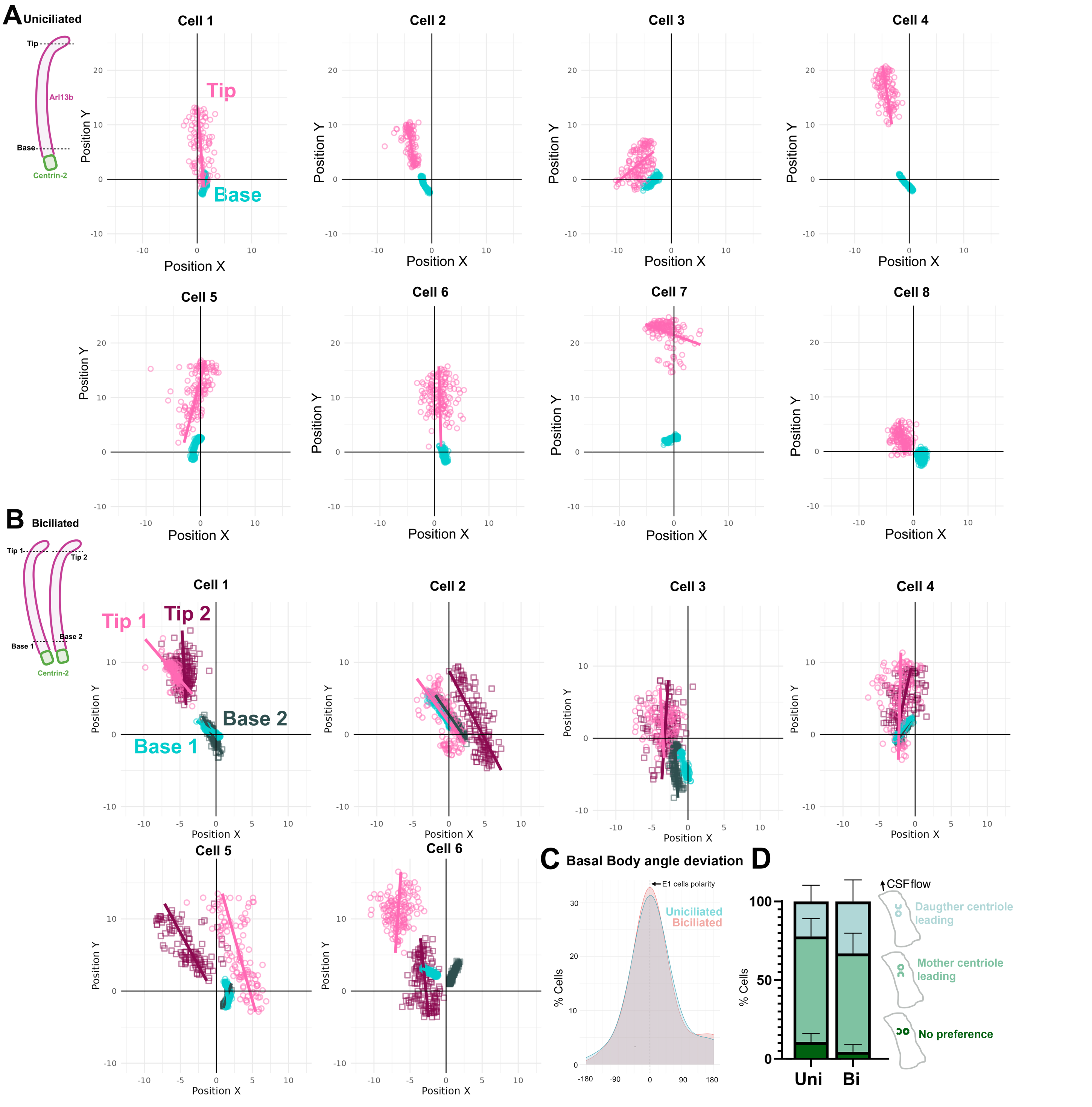


**Figure S3. E2 cells cilia movement and polarity.** Related to Figure 3. (**A, B**) Scattered plots of cilia position (frame-by-frame) of the tip (pink) and the proximal (green) segments of 8 uniciliated (A) and 6 biciliated (B) E2 cells. (**C**) Histogram of uni- and biciliated cells basal bodies angle deviation. **D**) Bar plots showing the proportion of E2 cells in which the mother centriole, the daughter centriole, or both centrioles at the same position were located at the leading edge of the basal body pair in the direction of translational polarity.

#### Supplementary Figure 4

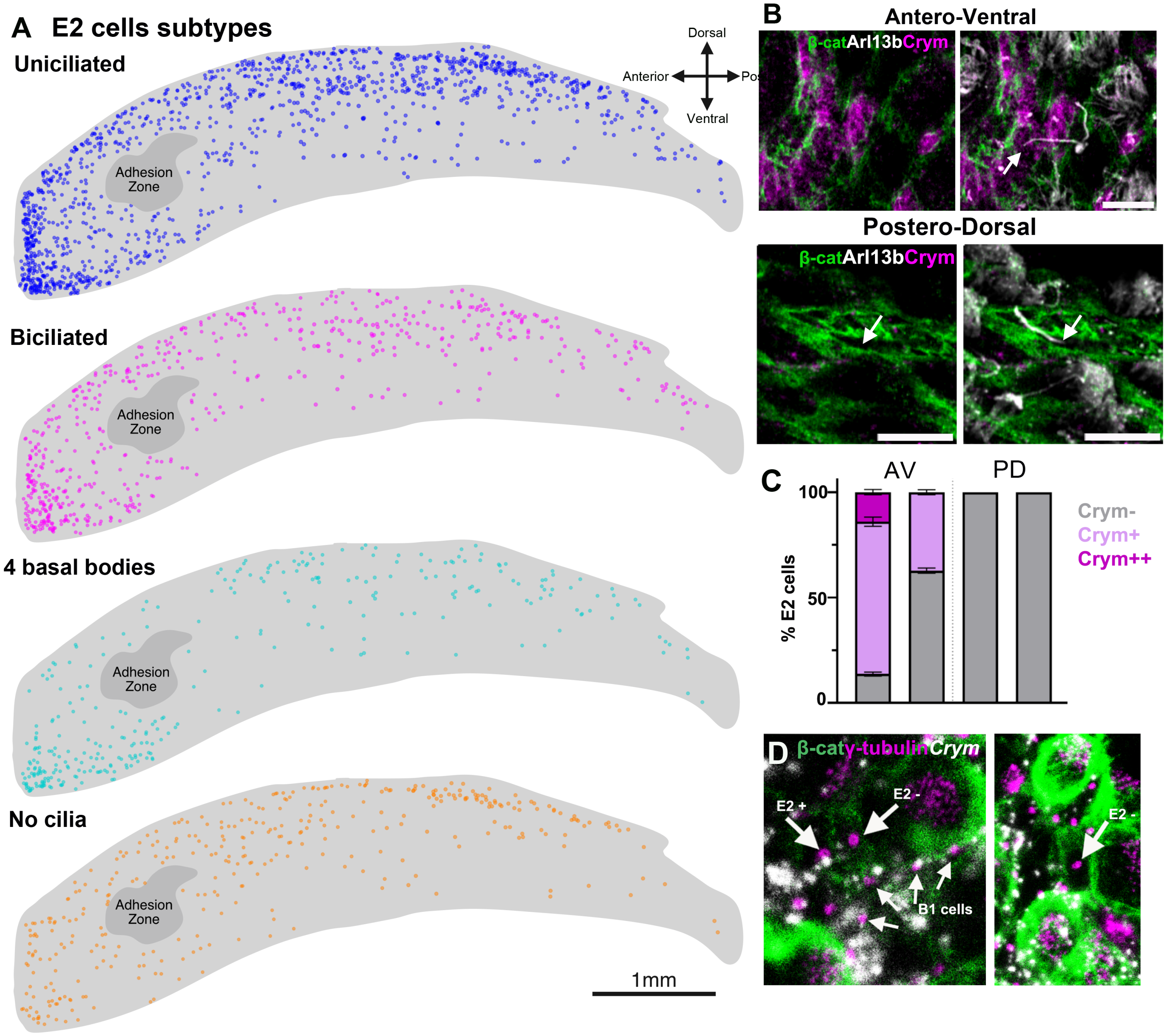


**Figure S4. Distribution of E2 cell subtypes in the lateral wall.** Related to Figure 4. **(A)** Maps of E2 cell subtypes distribution. The compass indicates anterior (A), posterior (P), ventral (V), and dorsal (D) orientation. (A) **B)** Confocal images of a whole mount of the lateral wall of the LV stained with Crym (magenta), b-catenin (green), and Arl13b (white). Note that PD E2 cells are negative for Crym. **C)** Barplots of the percentage of uniciliated and biciliated E2 cells Crym high (++), Crym low (+), and Crym- in the AV and PD regions. **D**) Confocal images of dual RNAscope–immunofluorescence in an en face section of the lateral ventricular wall, showing *Crym*$^{+}$ and *Crym*$^{-}$ E2 cells identified by γ-tubulin (magenta) and β-catenin (green) staining. Scale bars, (A) 2mm, (B) 10mm.

#### Supplementary Figure 5

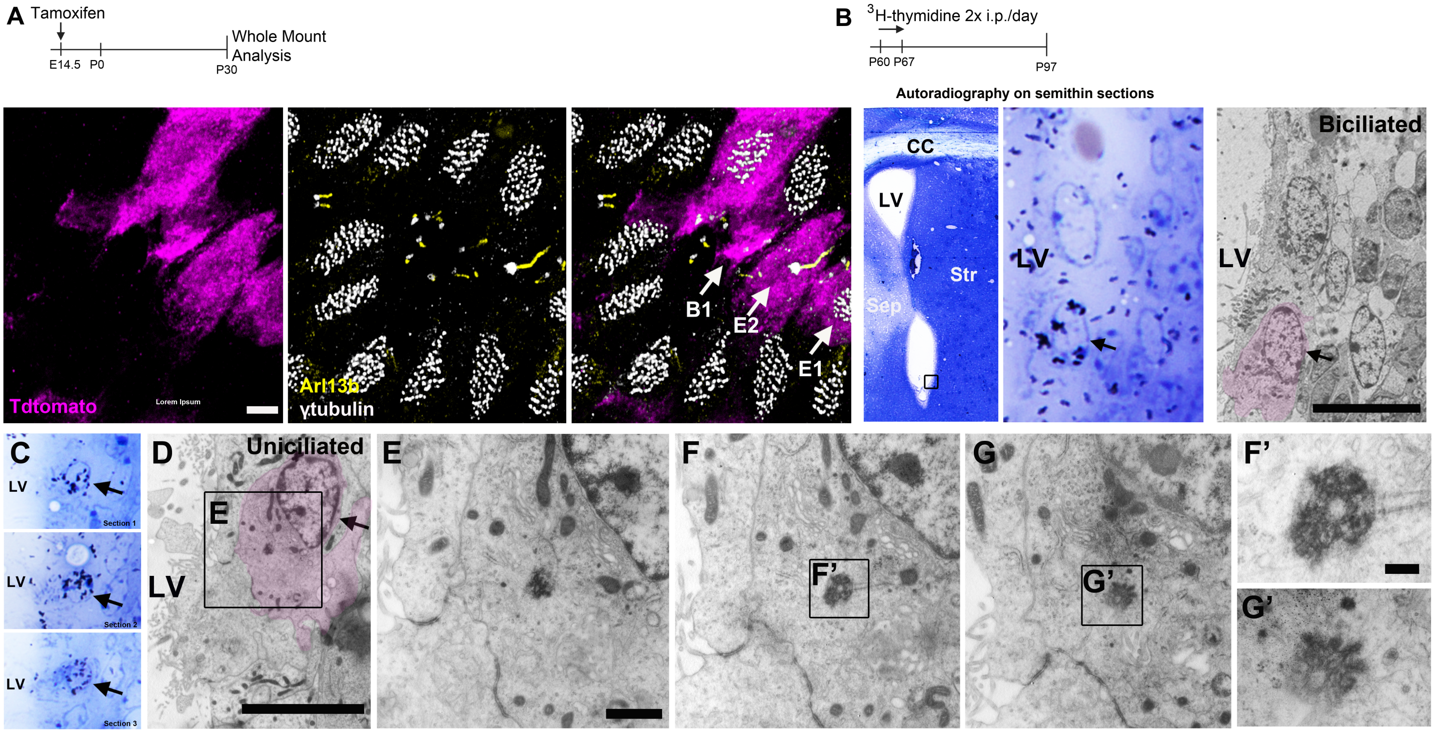


**Figure S5. E2 cells embryonic and postnatal origins.** Related to Figure 5. (**A**) Confocal images of Tdtomato+ clones containing E2, B1 and E1 cells. (**B**) Experimental outline: P60 mice received two i.p. injections of 3H-Thymidine (3H-Thy) /day for 7 days. The V-SVZ was analyzed one month after the last injection. Autoradiography of a toluidine blue-stained semithin section of the V-SVZ showing a $^3$H-thymidine+ label retaining cell (arrow) shown in Figure 5. (**C**) Autoradiography of 3 consecutive toluidine blue-stained semithin sections (1.5um) of the V-SVZ showing a $^3$H-thymidine+ label retaining cell (arrow). (**D-G**) TEM micrographs of the $^3$H-Thy-labeled cell in (C ) identified as a uniciliated E2 cell. Three serial ultrathin section images of the large electron dense basal body and their higher magnification images (F, G). Scale bars: 1 μm (D,E), 5μm (A, B, C) and 200nm (F, G).
